# CRISPR screens in iPSC-derived neurons reveal principles of tau proteostasis

**DOI:** 10.1101/2023.06.16.545386

**Authors:** Avi J. Samelson, Nabeela Ariqat, Justin McKetney, Gita Rohanitazangi, Celeste Parra Bravo, Rudra Bose, Kyle J. Travaglini, Victor L. Lam, Darrin Goodness, Gary Dixon, Emily Marzette, Julianne Jin, Ruilin Tian, Eric Tse, Romany Abskharon, Henry Pan, Emma C. Carroll, Rosalie E. Lawrence, Jason E. Gestwicki, Jessica E. Rexach, David Eisenberg, Nicholas M. Kanaan, Daniel R. Southworth, John D. Gross, Li Gan, Danielle L. Swaney, Martin Kampmann

## Abstract

Aggregation of the protein tau is a hallmark of Alzheimer’s disease and other tauopathies. Specific neuronal subtypes are selectively vulnerable to tau aggregation, but the underlying mechanisms are unknown. To systematically uncover the cellular factors controlling the accumulation of tau aggregates in human neurons, we conducted a genome-wide CRISPRi-based modifier screen in iPSC-derived neurons. The screen uncovered expected pathways, including autophagy, but also unexpected pathways, including UFMylation and GPI anchor synthesis, that control tau oligomer levels. We discover that the E3 ubiquitin ligase CUL5^SOCS4^ is a potent modifier of tau levels in human neurons, ubiquitinates tau, and is correlated with resilience to tauopathies in human disease. Disruption of mitochondrial function promotes proteasomal misprocessing of tau, which generates tau proteolytic fragments like those in disease and changes tau aggregation *in vitro*. These results reveal new principles of tau proteostasis in human neurons and pinpoint potential therapeutic targets for tauopathies.

## Introduction

Tauopathies, which include Alzheimer’s disease (AD), are a diverse group of neurodegenerative diseases defined by pathological aggregation of the protein tau and lack effective therapeutics. Tau is an intrinsically disordered protein and its canonical function is to bind to and regulate the stability and dynamics of microtubules. Recent work has established the role of tau in diverse neuronal processes, including axonal transport and synaptic transmission^1–3^. Mutations in the gene that encodes tau, *MAPT*, cause familial forms of frontotemporal lobar degeneration (FTLD)^4–6^. Most tauopathy cases, including in AD, however, are not caused by mutations in tau^6,7^, suggesting that factors in the cellular environment contribute to onset of tau pathology.

A key characteristic of tauopathies is selective vulnerability: specific regions of the brain and neuronal subtypes within them are vulnerable to specific tauopathies^8–12^. Furthermore, recent structures of the tau aggregate cores from patients have revealed disease-specific tau aggregate structures^13^, suggesting that determinants of tau conformation in different cellular environments may drive distinct disease outcomes. Post-translational modification is one link between the cellular environment and tau conformation and tau indeed is highly post-translationally modified. Specific post-translational modifications are associated with disease and are known to accelerate tau aggregation *in vitro*^14,15^.

A major challenge has been to identify how and which cellular factors control tau conformational changes that are on-pathway to aggregation. GWAS studies^16–19^ uncover modifiers of disease risk, but do not provide molecular mechanisms. Similarly, single-cell transcriptomics^9,20,21^, can describe the factors differentially expressed in vulnerable versus resilient neuronal subtypes, but these studies lack direct experiments to pinpoint how those factors causally control tau aggregation.

Experimental model systems enable the mechanistic dissection of factors controlling tau aggregation, but the most common model systems have limitations in terms of physiological relevance (such as tau over-expression in non-neuronal cell types) or are not amenable to high-throughput functional experiments that are required to gain comprehensive understanding.

Here, we use tau conformational-specific antibodies as probes to perform CRISPR-based screens in iPSC-derived human neurons^22^ harboring the FTD causing tau mutation, *MAPT* V337M^23,24^. While *MAPT* V337M tau causes FTD, it adopts the same fibril conformation as tau in AD^25^, suggesting that modulators of *MAPT* V337M aggregation could be relevant to both FTD and AD. We utilize this screening methodology to comprehensively identify modifiers of tau conformations and levels, uncovering both expected and unexpected factors.

We find a tau E3 ubiquitin ligase complex, CRL5^SOCS4^, and show that its expression is correlated with resilience to tau aggregation in a tauopathy mouse model and resilience to neuronal death in human AD and primary tauopathies. We also find that acute oxidative stress in neurons induces a proteasome-derived tau fragment that is secreted into the media. This fragment is positive for the AD biomarker NTA^26–28^, and changes the kinetics and quaternary structure of tau fibril formation *in vitro*. Our results highlight the power of neuron-specific screening approaches to reveal mechanisms of tau proteostasis.

## Results

### MAPT V337M neurons accumulate tau oligomers

Tau oligomers are a species of tau aggregate that precede fibril formation and are thought to be more toxic than tau fibrils^29–32^. Since oligomers are known to be transient and low abundance, we characterized native cell lysates of either *MAPT* WT or *MAPT* V337M iPSC-derived neurons (Figure 1A). Dot blots with neurons lysed in native conditions and fractionated based on detergent solubility revealed elevated levels of tau oligomers in the *MAPT* V337M neurons (Figure 1A, Figure S1). Most of the oligomer signal was in the PBS-soluble fraction and did not require detergents for solubilization. The T22 antibody had the highest specificity for the *MAPT* V337M neurons; *MAPT* WT neurons had the same T22 signal as *MAPT* knockdown neurons, indicating that this non-specific signal was not due to tau expression.

**Figure 1:**
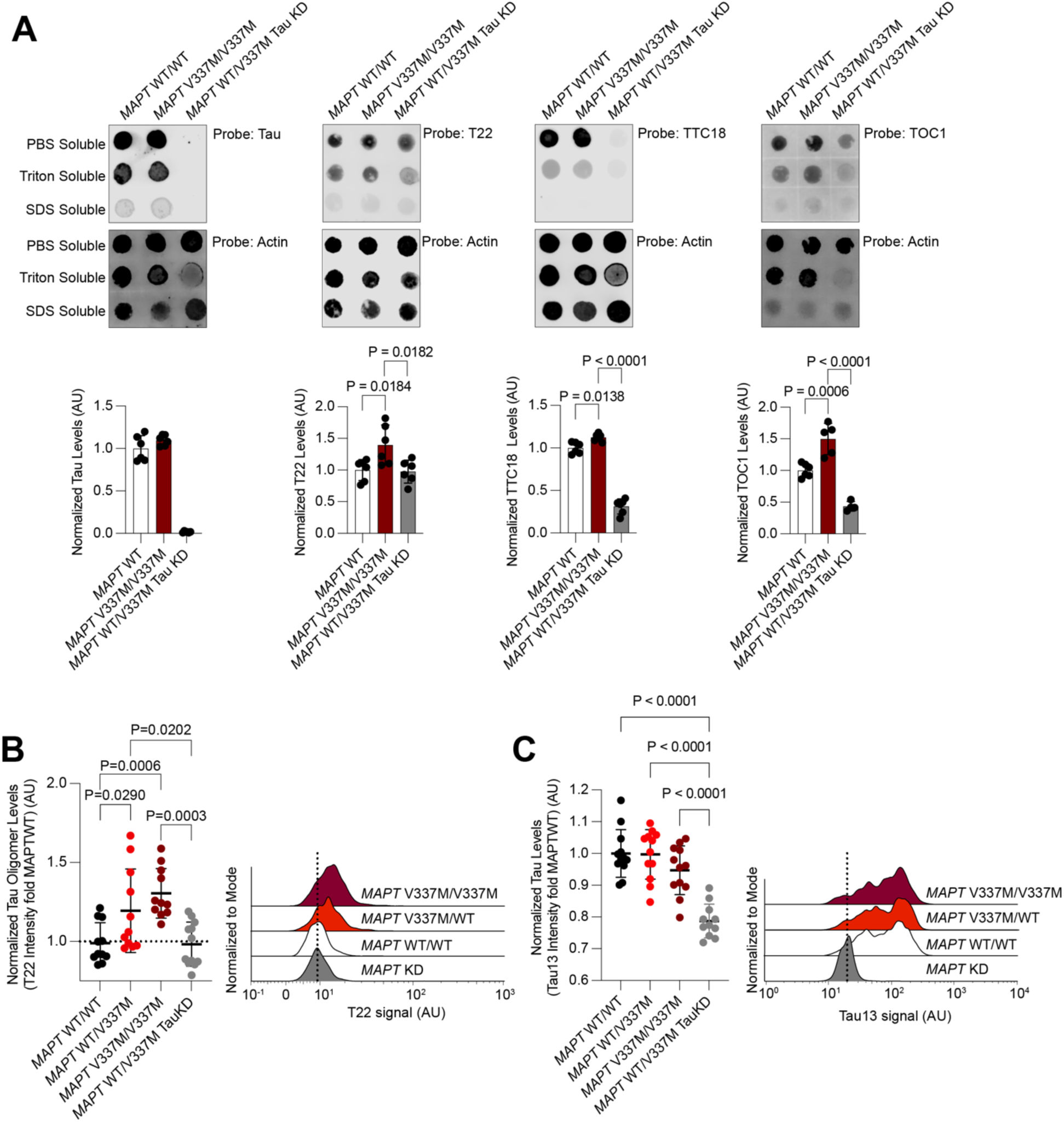
*MAPT V337M* neurons have higher levels of tau oligomers. **(A)** Dot blots (*above*) and quantification (*below*) of the PBS-soluble fraction of native neuronal lysates probed for total tau (tau13 antibody, *left)* or tau oligomers using different conformation-specific antibodies (T22, *center left*: TTC18, *center right*; TOC1, *right*). Neurons were lysed in DPBS and insoluble material was separated by centrifugation. Pellets were sequentially solubilized and then centrifuged in 0.1% Triton X-100 in DPBS and then 0.1% SDS in DPBS. All blots were normalized to actin (below) as a loading control. All samples are the average of six biological replicates except for TOC1, which is the average of four replicates, error bars are ±standard deviation. Blots used for quantitation are in Figure S1. **(B)** *Left,* Tau oligomer levels measured using the antibody T22 by flow cytometry in isogenic iPSC-derived neurons with one or two copies of the FTLD-causing *MAPT* V337M mutation. Intensities were normalized to the average of WT tau neurons (white). *Right,* Representative histograms are shown. **(C)** *Left,* Tau levels measured by flow cytometry using the total tau antibody tau13 in isogenic iPSC-derived neurons with one or two copies of the FTLD-causing *MAPT* V337M mutation. Intensities were normalized to the average of WT tau neurons (white). *Right,* representative histograms are shown. All samples are the average of twelve biological replicates, error bars are ±standard deviation. Standard one-way ANOVA was used for statistical analysis in **(A)-(C)**.

The T22 antibody is the only commercially available anti-tau oligomer antibody and has been used widely^29–31,33–35^. Therefore, we decided to use it as a probe for flow cytometry to enable genetic modifier screens based on T22 levels. Tau oligomer levels measured by flow cytometry recapitulated the dot blot data: *MAPT* V337M neurons had higher levels of T22 staining than *MAPT* WT neurons at 14 days post-differentiation (Figure 1B,C). Importantly, knockdown of *MAPT* using CRISPRi reduced levels of T22 staining (Figure 1B,C) confirming the dependence of T22 signal on tau expression and defining the tau-dependent vs. non-specific signal. Furthermore, we observed a dose-dependent effect of T22 staining dependent on the presence of one (*MAPT* WT/V337M) or two (*MAPT* V337M/V337M) copies of the V337M *MAPT* allele (Fig. 1B,C).

### Genome-wide screen for modifiers of tau oligomer levels

To identify cellular factors controlling tau oligomer levels in an unbiased genetic screen, we engineered the V337M iPSC line to express CRISPR interference (CRISPRi) machinery and optimized a protocol for the FACS sorting of fixed iPSC-derived neurons (see Methods). Using the anti-tau oligomer antibody T22 (Figure 2A), we conducted a genome-wide CRISPRi screen in two-week-old iPSC-derived neurons. Briefly, we infected iPSCs with our previously described next-generation lentiviral CRISPRi sgRNA library targeting all protein coding genes (five sgRNAs per gene, 104,535 sgRNAs) and 1,895 non-targeting controls^36^, differentiated them into neurons, and fixed them at Day 14 post-differentiation. Neurons were then stained with T22 antibody and an antibody against the neuronal marker NeuN. NeuN-positive cells were FACS-sorted into two bins: those neurons that had the highest and the lowest thirty percent of T22 signal. Frequencies of sgRNAs in each bin were determined by next-generation sequencing and compared to NTCs using a Mann-Whitney U test to assign p-values for each gene (Figure 2B, Supplemental Tables 2 and 3). 1,143 genes were called hits based on a false discovery rate (FDR) of 5% (see Methods). The knockdown of these hit genes causally modify T22 signal in iPSC-derived neurons.

**Figure 2:**
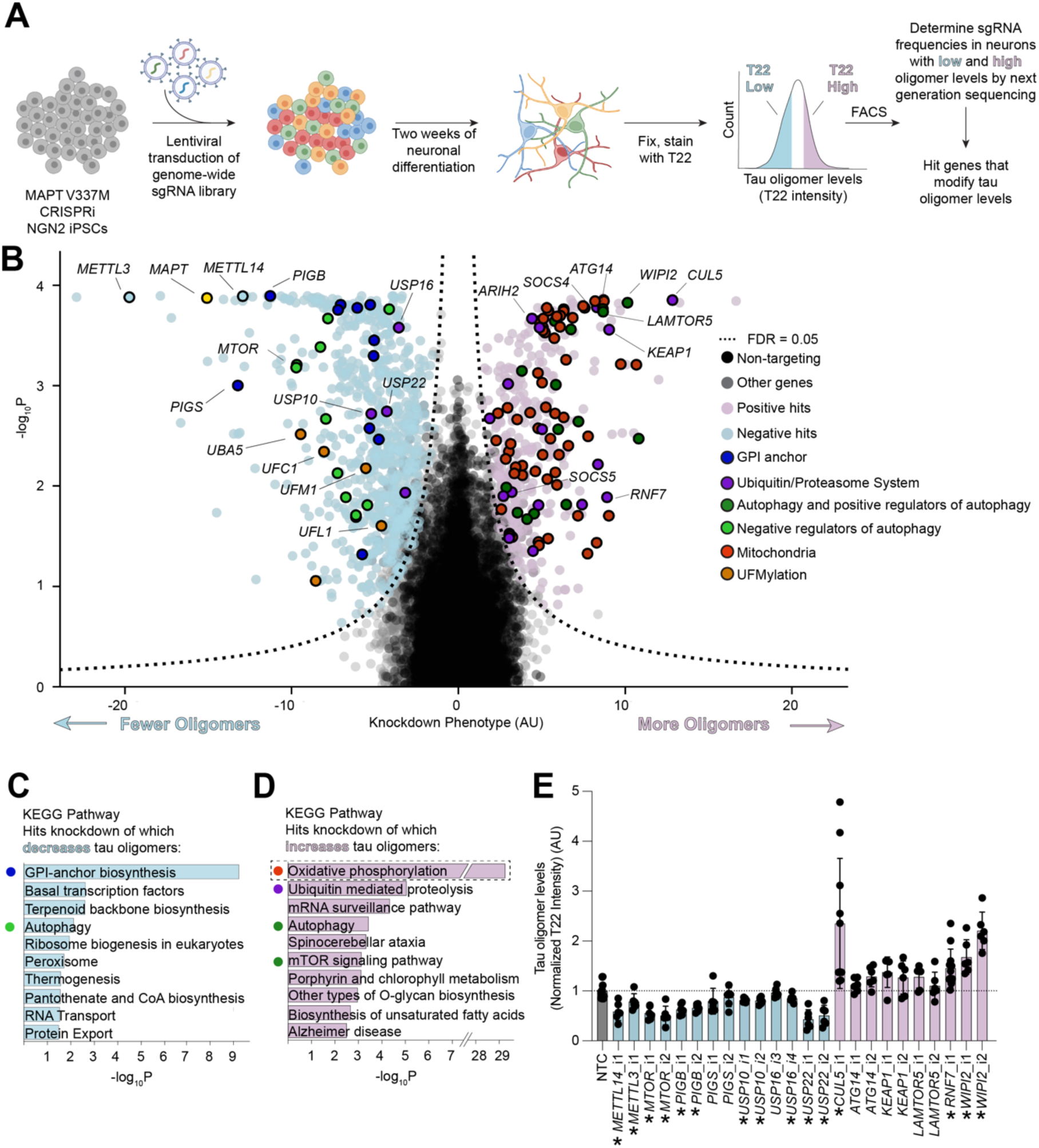
Genome-wide screen for tau oligomers levels in iPSC-derived neurons. **(A)** Schematic of screen. *MAPT* V337M heterozygous iPSCs were transduced with a pooled library of sgRNAs targeting every protein-coding gene. iPSCs were differentiated into excitatory neurons for two weeks, fixed, and stained with antibodies detecting the neuronal marker NeuN and tau oligomers (T22). The thirty percent of NeuN-positive cells with the lowest (blue) and highest (pink) tau oligomer signal were separated by FACS sorting. Genomic DNA was isolated from each population and the sgRNA cassette was sequenced using next-generation sequencing. Comparison of sgRNA frequencies was used to call hit genes. **(B)** Volcano plot of hit genes from genome-wide screen. Phenotype (normalized log_2_ ratio of counts in the T22-high versus T22-low populations), is plotted versus the negative log_10_ of the P-value, calculated with a Mann-Whitney U-test. Positive hits are in pink, and negative in light blue. Quasi-genes composed of random sets of non-targeting controls are in black and non-hit genes are in grey. *MAPT* is a top hit (yellow). (**C,D**) KEGG Pathway enrichment analysis uncovered oxidative phosphorylation and overlapping gene sets containing mitochondrial genes as the most significantly enriched term among genes knockdown of which increased tau oligomer levels (dashed box). To reveal additional pathways, a second round of enrichment analysis was performed after removal of mitochondrial genes. Top ten pathways by adjusted p-value are listed for (C) hits knockdown of which decreased tau oligomer levels (light blue) and for (D) hits knockdown of which increased tau oligomer levels (light pink). **(E)** Validation of select hit genes, labelled in (B), using individually cloned sgRNAs. Mean of six biological replicates is shown. Error bars are ±standard deviation. Bars in light blue were negative hits in the primary screen, and those in pink were positive hits. The asterisks denote sgRNAs with T22 levels significantly different from the non-targeting control, NTC (p <0.05 using a standard one-way ANOVA). sgRNAs *METTL14*i2 and *METTL3*i2 were toxic to iPSCs and excluded from this analysis.

KEGG pathway analysis of hit genes revealed several pathways and genes expected based on the literature, increasing confidence in the screen results (Figure 2B-D). *MAPT* was a top hit knockdown of which decreased tau oligomer levels, as expected. Known regulators of tau and tau oligomer levels, such as genes that control autophagy, were significantly enriched (Figure 2B-D). Knockdown of autophagy machinery and positive regulators of autophagy increased tau oligomer levels, and knockdown of negative regulators of autophagy such as *MTOR* decreased tau oligomer levels^37,37–40^. Knockdown of genes involved in the methyl-6- Adenosine (m^6^A) modification of RNA, including *METTL14* and *METTL3*, were among the top hits that decrease tau levels upon knockdown. A recent study has implicated m^6^A modification of RNA as a driver of tau-mediated neurodegeneration in mouse tauopathy models and human disease^41^. These results confirm that the methodology used here recapitulates shared characteristics of tau biology across systems.

Surprisingly, the strongest signature overall from the KEGG pathway analysis was oxidative phosphorylation, which includes genes encoding mitochondrial proteins, especially for the electron transport chain (ETC). Knockdown of these genes increases tau oligomer levels (Figure 2D). KEGG pathway analysis without mitochondrial hits (Figure 2D) revealed other pathways controlling tau oligomer levels. Autophagy and the Ubiquitin/Proteasome System (UPS) were significantly enriched, including the Cullin-RING ligase *CUL5*, and genes encoding four CUL5 interacting proteins, *SOCS4*, *SOCS5*, *ARIH2*, and *RNF7*. We identified GPI-anchor biosynthesis to be the most significantly enriched set of genes knockdown of which decreases tau oligomer levels (Figure 2C). We also identified an additional pathway that controls tau oligomerization, UFMylation, including the core UFMylation genes *UFC1*, *UFM1*, *UFL1*, and *UBA5*.

We then chose a subset of genes of 14 genes (26 sgRNAs) from the significantly enriched pathways to validate using individually cloned sgRNAs (Figure 2E). These experiments generally recapitulated the screen phenotypes.

### Secondary screens pinpoint cellular factors controlling tau levels and genotype-specific factors

To characterize the top hits more deeply, we cloned a small, pooled library targeting 1,037 hit genes (5 sgRNAs per gene) and 250 non-targeting controls (Supplemental Table 4). We then performed seven secondary CRISPRi screens using this library (Figure 3A; Supplemental Tables 2 and 3).

**Figure 3:**
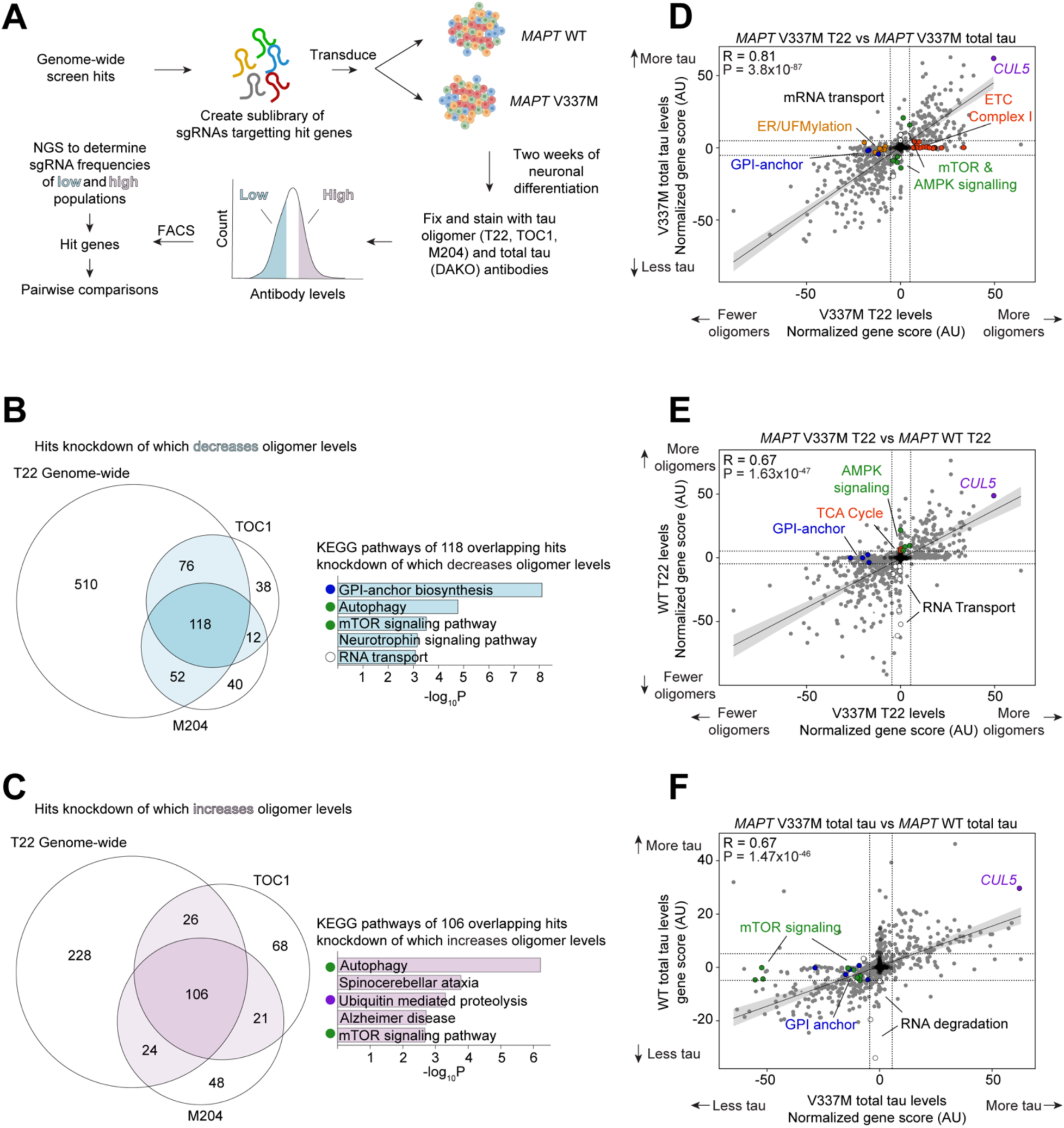
Secondary screens reveal modifiers specific to tau oligomer levels and *MAPT* genotype. **(A)** Schematic of retest strategy. A focused sgRNA library targeting all the hits from the genome-wide screen was screened in *MAPT* WT and *MAPT* V337M neurons using a panel of different antibodies. **(B)** Overlap of hit genes knockdown of which increases tau oligomer levels detected by three different tau oligomer antibodies, T22, TOC1, and M204. Top KEGG Pathways enriched for the 118 hit genes common to all three screens were calculated after removal of mitochondrial genes. **(C)** Overlap of hit genes knockdown of which decreases tau oligomer levels detected by three different tau oligomer antibodies, T22, TOC1, and M204. Top KEGG Pathways enriched for the hit genes common to all three screens are listed. **(D)** Comparison of total tau and tau oligomer screens in *MAPT* V337M neurons by gene score. Non targeting controls are in black and genes are in grey. ETC Complex I genes (red), ER/UFMylation genes (orange), and GPI-anchor genes (blue) were oligomer-specific hits. *CUL5* is in purple. mTOR and AMPK signaling genes are in green. **(E)** Comparison of tau oligomer screens in *MAPT* WT vs. V377M neurons by gene score. Non targeting controls are in black and genes are in grey. GPI-anchor genes (blue) were V337M-specific, while knockdown of genes involved in mRNA transport (white), especially nuclear pore subunits, strongly decreased tau oligomer levels in WT but not V337M neurons. Knockdown of TCA cycle genes (red) and genes involved in AMPK signaling (green) weakly increased tau oligomers in WT, but not V337M neurons. *CUL5* is in purple. **(F)** Non targeting controls are in black and genes are in grey. Comparison of screens for total tau levels in *MAPT* V337M versus WT neurons. Total tau levels in *MAPT* V337M neurons were more sensitive to knockdown of mTOR signaling (green) and GPI anchor genes (blue). Total tau levels in *MAPT* WT neurons were more sensitive to knockdown of genes involved in RNA degradation (white). *CUL5* is in purple.

Tau oligomer antibodies can have high levels of non-specific binding. To increase our confidence that genes from the primary screen truly control tau oligomer levels, we performed three additional screens: we used two alternative tau oligomer antibodies, TOC1^42^ and M204^43^, as well as a different lot of T22. There was a substantial overlap in modifiers of tau oligomers detected by all three antibodies (Figure 3B-C). Consistent with the primary screen, knockdown of genes involved in oxidative phosphorylation was the most enriched pathway that increased T22, TOC1, and M204 signal. KEGG Pathway analysis of the 108 shared hits without mitochondrial genes knockdown of which increased oligomer levels confirmed that autophagy, mTOR signaling, and the UPS are high-confidence pathways that control tau oligomer levels (Figure 3B-C). Conversely, knockdown of GPI-anchor biosynthesis and RNA transport genes decreased tau oligomer levels and were common to all three screens (Figure 3B).

To distinguish those hits that selectively control tau oligomer levels from those hits that control total tau levels, we performed a screen with the DAKO total tau antibody (Figure 3D). Comparison of gene scores between the two screens (see methods) revealed that knockdown of UFMylation, GPI-anchor, and electron transport chain genes all had much stronger effects on tau oligomer levels than on total tau levels. Knockdown of genes involved in the regulation of autophagy, such as mTOR, had stronger effects on total tau levels than on tau oligomers, consistent with the role of autophagy in clearing monomeric tau^37–40^.

To find hits unique to the *MAPT* V337M tau genotype, we next performed screens in a *MAPT* WT background using both T22 and total tau antibodies (Figure 3E-F). There was a broad overlap of the modifiers of total tau and tau oligomer levels in both genetic backgrounds, but some hits unique to each *MAPT* genotype. KEGG pathway analysis of genes unique to the WT tau line identified pathways involved in mRNA transport, while autophagy and mTOR signaling pathways were unique to the V337M line. Across all eight screens performed in this study, *CUL5*, a gene encoding an E3 ubiquitin ligase, was a top hit, regardless of *MAPT* genotype or antibody.

### CUL5 controls tau levels in neurons

CUL5 is a Cullin-RING E3 ubiquitin ligase that is best known for its role in regulating aspects of the immune response^44–47^. CUL5 serves as the scaffold for Cullin RING ligase complex 5 (CRL5) ,which facilitates ubiquitin transfer from E2 ubiquitin-conjugating enzymes to CRL5 substrates. RNF7, also a hit in the primary screen (see Figure 2B), is essential for binding of E2 enzymes to the CUL5 C-terminus. The E3 ligase ARIH2, also a hit in the primary screen (see Figure 2B), can form an E3-E3 complex with CUL5 and adds a priming round of monoubiquitin to the substrate in the absence of an E2 enzyme bound to the CRL5 complex^48–50^. The N-terminus of CUL5 binds to substrates via modular substrate adaptors containing a SOCS-box along with the proteins Elongin B and Elongin C (ELOB and ELOC). Activity of CRL5 is dependent on neddylation, a ubiquitin-like post-translational modification that is conjugated by the NEDD8-specific E2 enzyme UBE2F^51,52^.

Knockdown of both *CUL5* and *RNF7* with independently cloned sgRNAs recapitulated screen phenotypes by flow cytometry and Western blot (Figure 4A and B, Western blot in Figure S2A). Western blot showed smaller increases in tau levels upon *CUL5* and *RNF7* knockdown than flow cytometry. Flow cytometry samples are composed mostly of neuronal cell bodies, whereas Western blots samples contain the entire culture well of neurons. We hypothesized that tau levels may be differentially affected by *CUL5* and *RNF7* knockdown in the soma versus the neurites. Mechanical fractionation of neurons^53^ followed by Western blot revealed that CUL5 was localized primarily to the soma and that *CUL5* knockdown affected only tau levels in the soma (Figure 4C,D).

**Figure 4:**
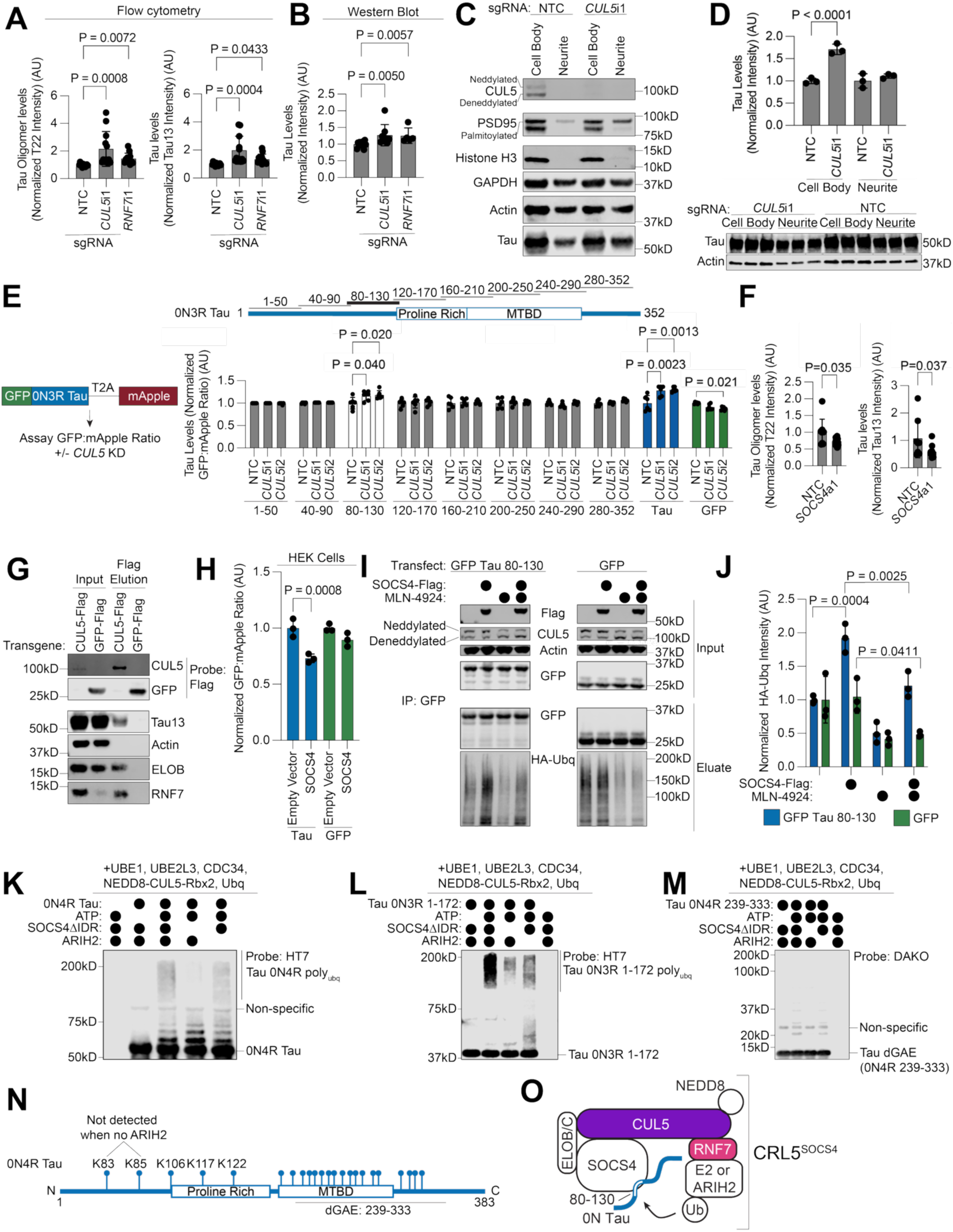
CRL5^SOCS4^ ubiquitinates tau and controls tau levels. **(A,B)** Individual knockdown of *CUL5* and *RNF7* reveals increases in Tau oligomer and total tau levels by flow cytometry (A) and western blot (B), relative to non-targeting control (NTC). Average of twelve biological replicates for flow cytometry, six biological replicates for Western blot (Western blots in Figure S2A). **(C)**. Mechanical fractionation of neurons into cell body and neurite fractions evaluated by western blot. CUL5 is only detected in the cell body fraction. Only mature, palmitoylated PSD95 is detected in the neurite fraction as expected. **(D)** *CUL5* KD increases tau levels in the cell body, but not neurite fraction. Quantitation (*top*) of Western blot data (*bottom*), tau levels normalized by actin loading control. Three biological replicates per sample. **(E)** Schematic of dual-fluorescence reporter for post-translational regulation of tau levels in neurons (*left*). Levels of each fifty-amino acid segment of tau sequence (schematic, *top*), tau (blue), and GFP (green) assayed in the presence of *CUL5* knockdown or non-targeting control sgRNA (*right*). Only tau 80-130 (white) recapitulates the *CUL5* knockdown sensitivity of full-length tau. Six biological replicates were used per sample. **(F)** Tau oligomer (*left*) and total tau levels (*right*) measured by flow cytometry following overexpression of SOCS4 using CRISPRa, compared to NTC. Average of nine biological replicates. **(G)** Immunoprecipitation of CUL5-Flag or GFP-flag reveals tau in the CUL5-flag eluate, along with ELOB and RNF7, markers of assembled CRL5 complexes. **(H)** Over-expression of SOCS4 in HEK cells with the dual-fluorescence reporter shown in (F) decreases tau levels (blue) but not GFP levels (green). Average of three biological replicates is shown. **(I)** Overexpression of SOCS4 in HEK cells increases the amount of ubiquitinated GFP-Tau 80-130 but not of ubiquitinated GFP. This increase in ubiquitination decreases upon treatment with the neddylation inhibitor, MLN4924. Gels representative of three biological replicates. **(J)** Quantitation of HA-ubiquitin signal in (I). Tau or GFP samples normalized to empty vector, vehicle treated samples. Three biological replicates. **(K-M)** *in vitro* ubiquitination reactions with three different substrates (K) 0N4R tau (L) 0N tau 1-172 (M) 0N4R Tau 239-333, also known as tau dGAE. Gels representative of three technical replicates. **(N)** Cartoon of ubiquitination sites identified by mass spectrometry of *in vitro* ubiquitination reactions labelled on equivalent sites on 0N4R tau. Lysines within the region 80-130 are labelled. MTBD, microtubule-binding domain. **(O)** Model of CRL5^SOCS4^-dependent ubiquitination of tau. For all applicable subpanels, one-way ANOVA was used for statistical analysis. P-values of >0.05 are not shown. Error bars are ±standard deviation.

To specifically measure the post-translational regulation of tau levels, we created a dual-fluorescence reporter for tau levels in neurons using a GFP-tau construct with a C-terminal T2A- mApple cassette (Figure 4E). Since both GFP and mApple are transcribed from the same transgene, the GFP-to-mApple ratio is a measure of post-translational changes in tau levels.

Knockdown of *CUL5* increased the GFP-to-mApple ratio only for GFP-tau and not for GFP alone (Figure 4E). The magnitude of increase of the reporter is similar to that of the soma-only western blots, within measurement error (Figure 4D). Treatment of neurons with the proteasome inhibitor MG132 reversed the effect of *CUL5* knockdown on the GFP-to-mApple ratio, suggesting that CRL5-dependent clearance of tau is proteasome-dependent (Figure S2B).

Cullin-ring ligases like CRL5 recognize substrates via short amino acid sequences called degrons. To find the region of tau that acts as a CRL5 degron, we cloned sliding windows of fifty amino acids of the 0N3R tau sequence with a ten amino acid overlap into the dual-fluorescence reporter construct (Figure 4E). We made stable iPSC lines expressing these fifty amino acid segments using lentivirus, transduced them with sgRNAs against *CUL5* or non-targeting controls, differentiated them into neurons and assayed the GFP-to-mApple ratio 14 days post-differentiation. Along with full length tau, only the construct encompassing 0N3R tau residues 80-130 (2N4R tau residues 138-188), showed an increased GFP-to-mApple ratio upon *CUL5* knockdown. This increase in was dependent on proteasome activity, since it was abrogated by the proteasome inhibitor MG132 (Figure S2C).

CRLs bind substrates via substrate adaptor proteins that bind to the N-terminus of the CRL. In our genome-wide screen, two CRL5 adaptors were positive hits, *SOCS4* and *SOCS5* (Figure 2B, Figure S2D). We pursued *SOCS4* because it was a much stronger hit than *SOCS5*. Over-expression of *SOCS4* in neurons using CRISPRa decreased tau oligomer and total tau levels (Figure 4F). Because *CUL5* knockdown controls tau levels in a proteasome-dependent manner, and *SOCS4* overexpression decreases tau levels, we hypothesized that CRL5^SOCS4^ could be directly ubiquitinating tau in cells.

### CRL5^SOCS4^ ubiquitinates tau

To determine if tau and CUL5 physically interact, we overexpressed a 3xFlag-tagged CUL5 construct in iPSC-derived neurons. CUL5-FLAG, but not GFP-FLAG, co-immunoprecipitated with endogenous tau (Figure 4G). Importantly, ELOB and RNF7 were both present in the eluate, confirming that functional CUL5-3xFlag complexes were immunoprecipitated. Thus, CUL5 and tau physically interact.

We next sought to reconstitute tau-CRL5^SOCS4^ interactions in HEK cells and *in vitro*. SOCS4 overexpression in HEK cells decreased tau levels, but not GFP levels, as measured by the same tau dual-fluorescence reporter used above (Figure 4H). Tau is ubiquitinated on its C-terminus primarily by the E3 ubiquitin ligase CHIP in HEK cells^54–56^. To decouple SOCS4 ubiquitination from CHIP ubiquitination, we co-transfected HEK cells with an HA-tagged ubiquitin, SOCS4, and GFP-tau 80-130, which contains the *CUL5* knockdown-sensitive degron (Figure 4E). More ubiquitinated GFP-tau 80-130 was detected by GFP immunoprecipitation from HEK cells upon SOCS4 transfection than transfection with empty vector (Figure 4I-J).

Ubiquitination was not due to the GFP tag, as this SOCS4-dependent ubiquitination increase was not observed upon transfection with GFP alone (Figure 4I-J). The SOCS4-dependent increase in ubiquitination was reduced by the neddylation inhibitor MLN4924, confirming that SOCS4- dependent ubiquitination of tau requires neddylation (Figure 4I-J).

We next reconstituted the CRL5^SOCS4^ ubiquitination cascade *in vitro* with recombinant components purified from *E. coli*. We co-expressed ELOB and ELOC with either full-length SOCS4 or a construct that omits the SOCS4 intrinsically disordered domain (IDR), SOCS4ΔIDR^57^. By size exclusion chromatography, full-length SOCS4-ELOBC eluted in the void volume of a S200 column, suggesting improperly folded or aggregated SOCS4 (Figure S2E-F). Therefore, we proceeded with the SOCS4ΔIDR construct. Accumulation of poly-ubiquitinated tau was dependent on SOCS4ΔIDR and ATP (Figure 4K). Tau conjugated to 1, 2, 3, or 4 ubiquitins accumulated in the absence of SOCS4ΔIDR, suggesting non-specific interactions of tau with CUL5 and/or the E3 ligase ARIH2. Poly-ubiquitinated tau as well as tau conjugated to 1-4 ubiquitins was decreased in the absence of ARIH2. In agreement with our results mapping the *CUL5*-dependent degron to residues 80-130, ubiquitination of 0N3R tau 1-172 was also SOCS4ΔIDR dependent (Figure 4L) but not tau dGAE (0N4R tau 239-333) (Figure 4M). The lack of conjugates of 4 or less ubiquitins to the 0N3R tau 1-172 construct suggests that the C-terminus of tau is responsible for SOCS4ΔIDR-independent ubiquitination products (Figure 4L). We then mapped ubiquitination sites by mass spectrometry (Figure 4N). Overall, there were 23 total ubiquitination sites mapped (Supplemental Table 5), with five sites identified between residues 80-130: K83, K85, K106, K117, and K122 (Figure 4N). Of these, two were not identified when ARIH2 was omitted from the *in vitro* ubiquitination reaction, K83 and K85. These data suggest a model where CRL5^SOCS4^ can extend poly-ubiquitin chains on the lysines between tau resides 80-130 (Figure 4O).

### CUL5 expression is correlated with disease phenotypes in human tauopathies

Given our finding that CRL5^SOCS4^ ubiquitinates tau for proteasomal degradation in cells and *in vitro*, we next asked whether CRL5^SOCS4^ may be a determinant of neuronal resilience to tauopathy in human patients. Analysis of snRNA-seq from human brain previously published from our lab^9^ found that *CUL5* is expressed more highly in human entorhinal cortex neurons resilient to AD than in those that are selectively vulnerable (Figure 5A).

**Figure 5:**
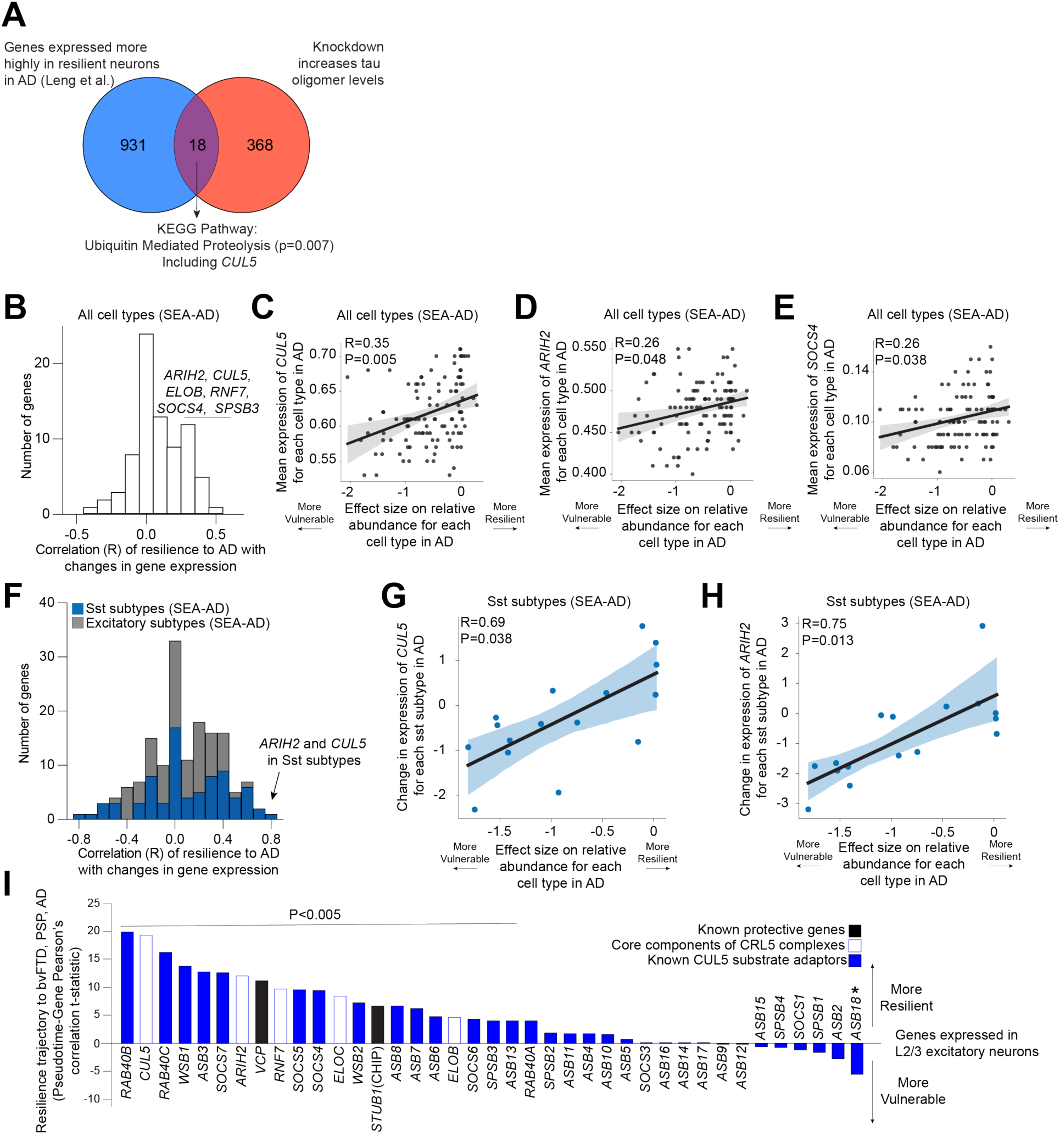
*CUL5* expression is correlated with lower vulnerability to tau aggregation in mice and to neuronal death in human Alzheimer’s Disease (AD). **(A)** Overlap between genes expressed more highly in excitatory neurons in the human entorhinal cortex that are resilient versus vulnerable to AD^9^ and genes that increase tau oligomer levels in this study. **(B)** Correlation of mean expression or change in expression of genes of known CRL5 components including *CUL5* with vulnerability of all cell types in AD. *ARIH2*, *CUL5*, *SOCS4*, *ELOB*, *RNF7* and *SPSB3* are all within the top three histogram bins. **(C)**-**(E)** Correlation of mean expression of (C) *CUL5* (D) *ARIH2*, and (E) *SOCS4* with vulnerability of all brain cell types in AD. **(F)** Correlation of mean expression and change in expression of genes of known CRL5 components including *CUL5* with vulnerability of excitatory (grey) and Somatostatin (Sst, blue) neuronal subtypes in AD. *ARIH2* and *CUL5* are within the top histogram bin. **(G)**-**(H)** Correlation of change in expression of (G) *CUL5* and (H) *ARIH2* with vulnerability of Sst neuronal subtypes in AD. For C-D and G-H, error bars represent the 95% confidence interval over 1,000 bootstraps. p-value is the Pearson correlation p-value corrected for multiple-hypothesis testing using the Benjamini-Hochberg procedure. **(I)** Gene expression correlation with the relative trajectory of neuronal resilience (positive) versus vulnerability (negative) in layer 2/3 IT neurons, based on Rexach et al^12^. Predicted effects on resilience vs. vulnerability based on gene expression patterns in human bvFTD, AD, and PSP brain tissues (y-axis, Pearson’s correlation t-statistic). All core proteins of CRL5 complexes (white with blue outline), all known CUL5 substrate adaptors (blue) and members of the ubiquitin/proteasome system with established protective roles in tauopathies (black). Two-sided Pearson’s correlation with FDR correction across 30,309 transcripts, n = 12,218 cells; all genes shown have FDR < 0.05. p-value is the Pearson correlation p-value adjusted with FDR correction across the numbers of genes as in Rexach et al^12^. Asterisk denotes p<0.005.

We next analyzed the Seattle Alzheimer’s Disease Brain Cell Atlas^58^ to determine if genes involved in CRL5-dependent ubiquitination are correlated with cell loss in AD (Figure 5B, Supplemental Table 6). Across all cell types, higher *CUL5*, *RNF7, ARIH2*, and *SOCS4* expression are weakly but significantly correlated with resilience to cell death in AD (Figure 5C-E, Supplemental Table 6). The CRL5 substrate adaptor SPSB3, which was recently shown to be responsible for degrading nuclear cGAS/STING^47^, was also significantly correlated with resilience to cell death in disease. A more granular analysis of genes involved in CRL5- dependent ubiquitination in excitatory neurons and Somatostatin (Sst) inhibitory neurons revealed that decreases in *CUL5* and *ARIH2* expression are highly and significantly correlated with loss of vulnerable Sst inhibitory neurons in AD (Figure 5F-H).

To see if expression of *CUL5* and its interactors is correlated with resilience in other tauopathies, we analyzed a single cell multi-omic dataset^12^ of AD, bvFTD, and PSP. We compared gene expression changes along a pseudotime trajectory that proceeds from L2/3 IT neurons depleted in bvFTD (vulnerable) to resilient L2/3 IT neurons in AD, bvFTD, and PSP in the insular cortex. We found that expression of *CUL5*, *ARIH2*, *SOCS4*, and several other CUL5 interactors was significantly correlated with the resilient trajectory (Figure 5I). Expression of *CUL5*, *ARIH2*, and *SOCS4* are more highly correlated with resilience in this pseudotime trajectory than genes encoding other members of the UPS system known to play a role in tau proteostasis regulation, *VCP*^59,60^ and *STUB1*^61,62^ (the gene that encodes for CHIP).

These correlative findings, in conjunction with our biochemical results, suggest that CUL5, and proteins involved in CRL5-dependent ubiquitination, are determinants of neuronal vulnerability and resilience in tauopathies.

### Electron transport chain inhibition leads to the generation of a 25kD N-terminal tau fragment

Genes essential for the function of mitochondria, especially those encoding components of the electron transport chain (ETC), were the largest and most significant class of hit genes knockdown of which increased tau and tau oligomer levels in our primary and secondary screens. To validate these hits using an orthogonal approach, we investigated the effect of pharmacological ETC inhibition on tau oligomer levels.

We measured tau and T22 levels by flow cytometry after treating neurons for 24 hours with increasing concentrations of rotenone, an ETC complex I inhibitor (Figure 6A-B). In *MAPT* V337M neurons, rotenone treatment increased both total tau and tau oligomer levels in a dose-dependent manner, whereas in *MAPT* WT neurons, rotenone increased only total tau levels, but not tau oligomer levels. Validation of the increase in tau levels by Western blot revealed that rotenone promoted the formation of a ∼25kD tau fragment detected by the Tau13 antibody, which recognizes tau residues 13-21^63^ (Figure 6C). Fragment formation was independent of *MAPT* genotype, as treatment with rotenone also induced this fragment in *MAPT* WT neurons (Figure S3A). Surprisingly, we did not observe the complementary C-terminal fragment, suggesting that this fragment is degraded (Figure S3C). Over-expression of a GFP-0N3R tau transgene revealed the same molecular weight shift after rotenone treatment (Figure 6D), confirming that this fragment is due to post-transcriptional processing of tau.

**Figure 6:**
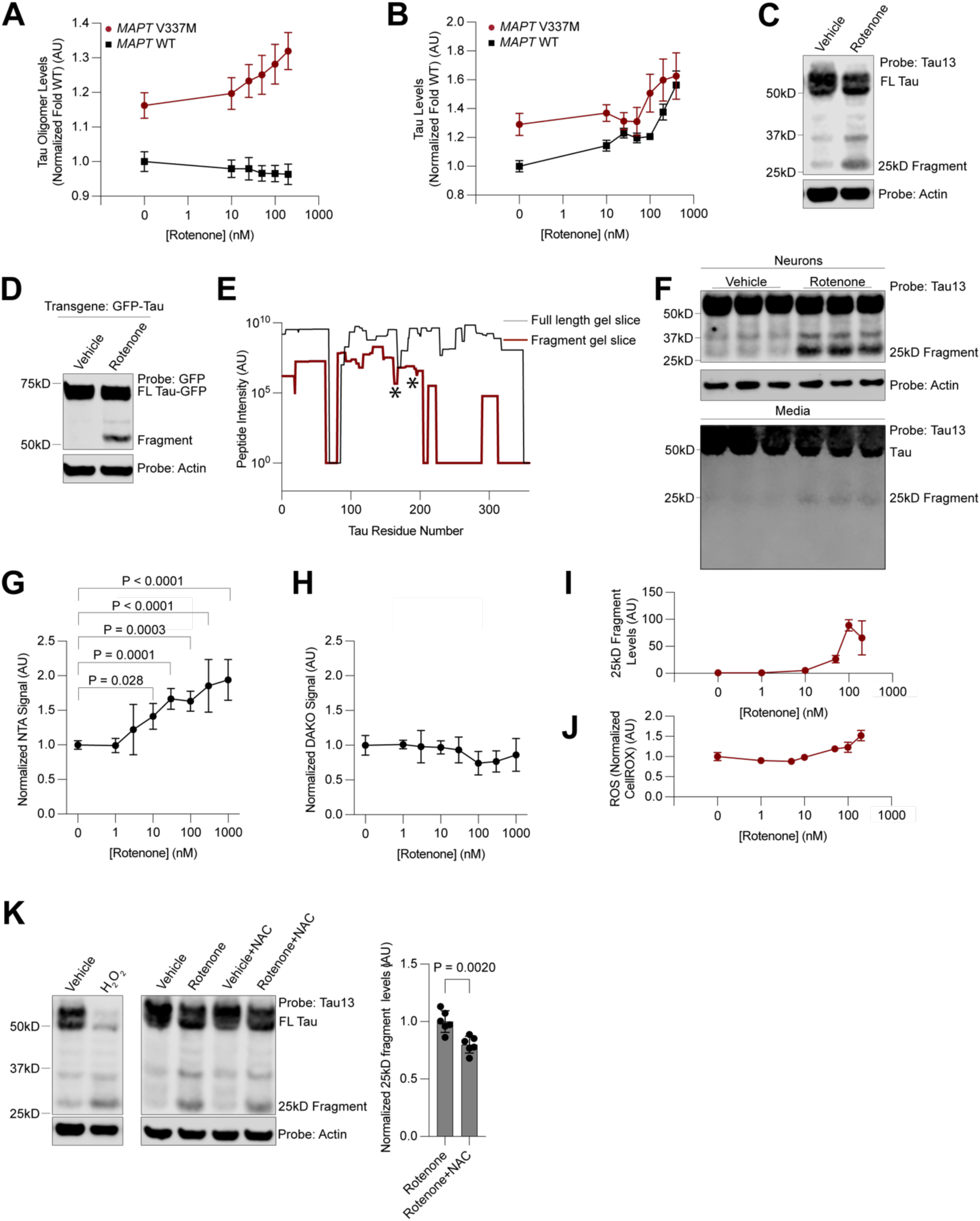
Rotenone increases ROS, tau oligomer, and total tau levels and induces a tau 25kD fragment. **(A,B)** Rotenone treatment increases levels of tau oligomers (A) and total tau (B) in *MAPT* V337M neurons, but only total tau levels in *MAPT* WT neurons in a dose-dependent manner. No significance test was performed. **(C)** Western blot of rotenone-treated neurons reveals a tau cleavage fragment. **(D)** Rotenone treatment of neurons results in an equivalent fragment from a GFP-Tau transgene. **(E)** Traces of averaged intensity from peptides per amino acid derived from mass spectrometry data for full-length tau (black) and rotenone induced fragment (dark red). A sharp decrease in intensity is seen in the rotenone induced fragment gel slice after residue 200, except for one peptide (see Figure S6). Stars represent neo-tryptic termini identified upon digestion with GluC, pinpointing two possible C-termini for the fragment, narrowing the fragment identity to residues 172-200 (2N4R numbering: 230-258). **(F)** Western blot of cell lysate (top) and undiluted conditioned media (bottom) shows the presence of the 25 kD fragment in the media. **(G,H)** ELISA signal for (G) NTA and (H) DAKO-based ELISA as a function of rotenone concentration. **(I)** Neuronal levels of the 25 kD fragment quantified by Western blot as a function of rotenone concentration. **(J)** ROS levels quantified by CellRox and flow cytometry as a function of rotenone concentration. **(K)** Treatment with 100µM hydrogen peroxide (*left*) increases 25 kD fragment formation, while treatment with the antioxidant N-Acetyl-Cysteine at 1µM (*middle*) decreases fragment formation (quantitation *right*). If not otherwise shown, rotenone concentration is 200 nM and treatment time is 24 hours. In applicable panels, one-way ANOVA was used for all statistical analysis. Unless otherwise noted, all samples are the average of six biological replicates, error bars are ±standard deviation. P-values >0.05 are not shown.

We then over-expressed GFP-Tau in iPSC-derived neurons and purified the rotenone-induced fragment for identification by mass spectrometry (Figure 6E). We observed a substantial decrease in the presence and intensity of peptides at approximately the 200th residue of the 0N3R tau sequence. However, considering that we did not identify any non-tryptic peptides in that region that would serve as the C-terminus for a fragment digested by an endogenous protease, we hypothesized that the digestion may be performed by a protease with trypsin-like cleavage site preference, such as the proteasome. This led us to pursue a second strategy, an in-solution digest of GFP-tau purified from rotenone-treated neurons using the non-tryptic protease GluC. We identified three peptides that were semi-specific for GluC with tryptic N termini, ending at 172 and 176 in the 0N3R MAPT sequence. Thus, the fragment sequence is narrowed to a small region spanning residues 172-200 in this region.

This fragment sequence is remarkably similar to the NTA tau biomarker^26–28,64^, a highly accurate biomarker in the CSF and plasma for NFT burden and cognitive decline in AD. We therefore hypothesized that the 25kD tau fragment may be released into the extracellular space. Indeed, we detected the presence of the 25kD tau fragment, but not tau fragments of other molecular weights, in the conditioned media of neurons treated with rotenone by Western blot (Figure 6F). We next performed the NTA ELISA and a C-terminal tau ELISA on neuronal media after rotenone treatment. Only the NTA ELISA, but not the C-terminal tau ELISA, showed an increase in signal as a function of rotenone concentration (Figure 6G-H). Therefore, we conclude that rotenone treatment specifically leads to the release of NTA-positive tau species and not general release of tau into the media.

### Acute oxidative stress leads to the generation of the 25kD tau fragment

Our use of iPSC-derived neurons allows us to determine the mechanism by which the 25kD tau fragment is derived and gain insight into a possible alternative mechanism by which NTA-positive tau fragments are formed. Treatment with Antimycin A, an ETC complex III inhibitor, but not CCCP, a proton-gradient uncoupler, or Oligomycin, an ATP Synthase inhibitor, also led to 25kD fragment formation (Figure S3D). ETC complexes I and III are generally considered the major sources of mitochondrial reactive oxygen species (ROS). Therefore, we hypothesized that ROS accumulation leads to 25kD fragment formation.

Indeed, increasing concentrations of rotenone, which led to a dose-dependent increase in the formation of the 25 kD fragment (Figure 6I) and mirrored increased in tau and tau oligomer levels, (Figure 6A,B) also led to a concomitant increase in neuronal ROS levels measured using the CellRox dye (Figure 6J). Neurons treated with hydrogen peroxide revealed close to complete conversion of full-length tau into the 25kD fragment (Figure 6K). The antioxidant N-acetyl-cysteine partially counteracted the rotenone-induced formation of the 25kD fragment (Figure 6K). 25kD fragment formation was not due to cell death via apoptosis or ferroptosis, as treatment with the pan-caspase inhibitor ZVAD-MVK, the ferroptosis inhibitor ferrostatin, or the ferroptosis inducer RSL3 did not change fragment levels (Figure S3E-H). Thus, acute increases in ROS levels are sufficient to generate the 25kD fragment.

To ask whether a broader set of factors that affect neuronal ROS levels also affect tau oligomer levels, we leveraged our previously published genome-wide screen for modifiers of neuronal ROS levels^65^. Systematic comparison of the knockdown phenotypes of all genes on ROS levels versus tau oligomer levels uncovered groups of genes that affected both in the same direction (Figure S4A). Knockdown of ETC components and other mitochondrial proteins such as *FECH* and *FH*, regulators of the oxidative stress response (*KEAP1*), and autophagy-lysosome factors (*WIPI2*, *CTSD*, and *PSAP*) increased both ROS and tau oligomers, whereas knockdown of GPI anchor genes and mTOR signaling factors decreased both ROS and tau oligomers (Figure S4A).

### Oxidative stress induced changes to proteasome activity lead to 25kD fragment accumulation

To identify the cellular protease that induces the 25kD fragment, we treated neurons with rotenone and inhibitors of common tau proteases: cathepsins, calpain, and the proteasome^66–69^ (Figure S3I-J). Only inhibition of the proteasome decreased 25kD fragment formation (Figure 7A). Proteasome activity measured by native gel revealed a fifty percent decrease in activity of 30S, 26S and 20S proteasomes upon rotenone treatment, although no changes in levels of proteasomal proteins were observed (Figure 7B and C, Figure S3K). Proteasomes isolated from synaptosomes showed the same changes in activity as the overall pool of proteasomes (Figure S3L-N). Similarly, treatment with a membrane-impermeable proteasome inhibitor ruled out the involvement of membrane-bound proteasomes in the production or extracellular release of the 25kD fragment (Figure S3O). Thus, we hypothesize that changes in proteasome processivity, rather than levels, may promote formation of the 25kD fragment.

**Figure 7:**
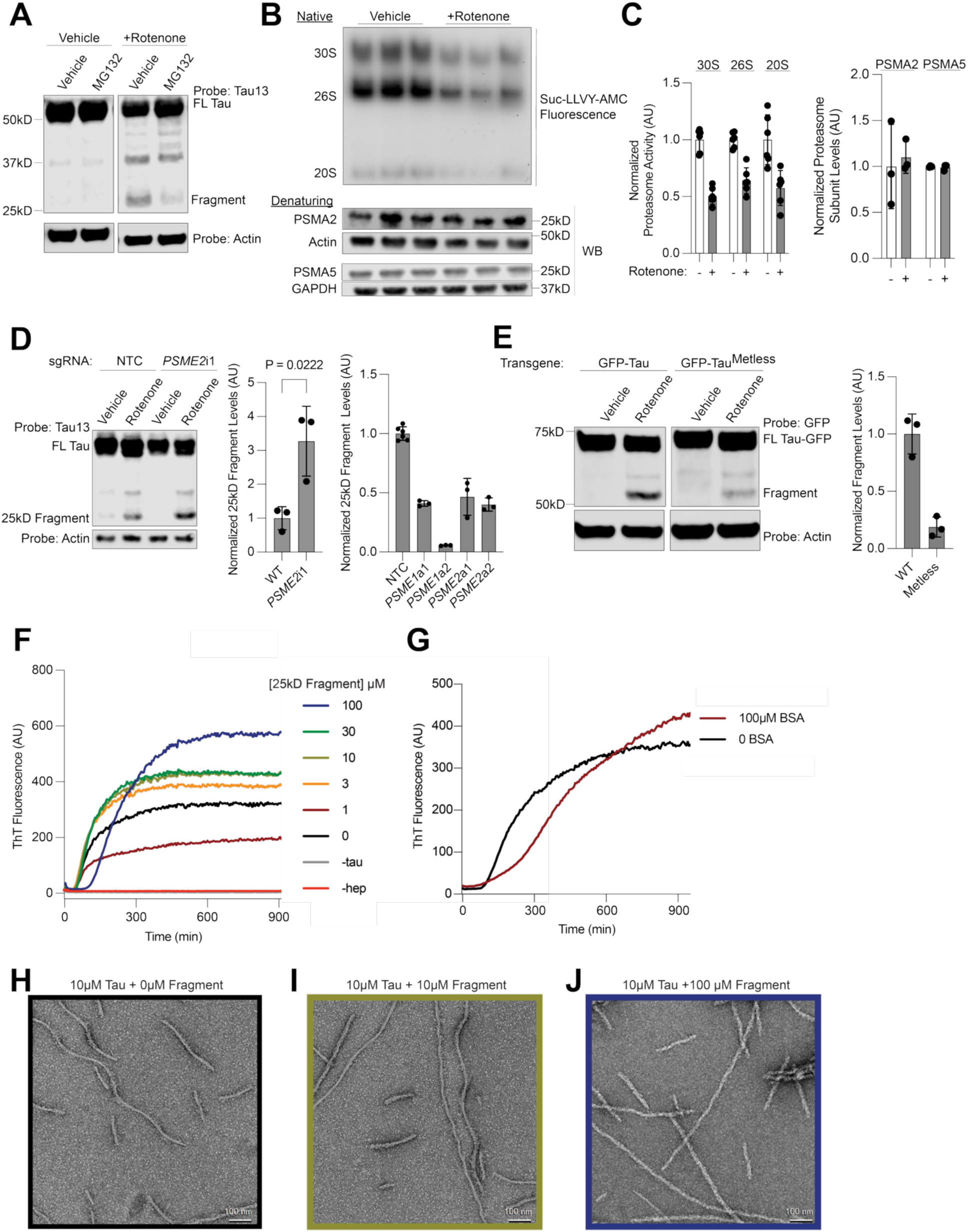
ROS leads to proteasome dysfunction which causes accumulation of the tau 25kD fragment. **(A)** Treatment with proteasome inhibitor MG132 at 1 µM decreases levels of the 25kD fragment. **(B)** Treatment with rotenone decreases proteasome activity (*top*) but not levels (*bottom*). **(C)** Quantitation of **(B)**. **(D)** Knockdown of the proteasome regulator *PSME2* increases levels of the 25 kD fragment (*left*) Western blot with quantitation. CRISPRa-mediated over-expression of *PSME1* or *PSME2* decreases fragment levels (*right*). Full gels in Figure S3P-Q. **(E)** Expression of a GFP-tau construct in which all methionines have been replaced by leucines (GFP-tau^Metless^) decreases fragment formation as compared to GFP-tau in neurons. Western blot (*left*), quantitation of western blot (*right*). **(F-G)** Increasing concentrations of 25kD fragment (F) but not BSA (G) increase the final ThT fluorescence of *in vitro* aggregation reactions of 10 µM 0N4R tau. Representative ThT traces of six technical replicates are shown. (**H-J)** Negative stain EM images of fibrils aggregated in the absence (H), or presence of 10 µM (I) or 100 µM (J) 25 kD fragment. Note straightening of fibrils in (H) and (J). Average of three biological replicates unless otherwise stated. One-way ANOVA used otherwise stated. Error bars are ±standard deviation.

Upon induction of oxidative stress, the proteasome activator PA28 is often up-regulated or increases its association with the 20S proteasome in order to deal with increased load of oxidized proteins^70,71^. Overexpression of the two PA28 subunits, PA28⍺ and PA28β, encoded by the genes *PSME1* and *PSME2* respectively, by CRISPRa, decreased 25kD fragment formation (Figure 7D, left). Knockdown of the PA28 subunit PA28β increased 25kD fragment levels (Figure 7C, right, Figure S3P-Q). Therefore, changes in proteasome activity, either general decreases in proteasome processivity caused by ROS or changes in PA28 occupancy can alter the formation of tau 25kD fragment levels.

We next asked whether direct oxidation of tau may also contribute to 25kD fragment formation. 0N3R tau only has one cysteine, C291, which is located close to the microtubule-binding domain. Expression of a GFP-0N3R tau transgene with the cysteine mutated to alanine (C291A) did not change fragment levels compared to a transgene with the cysteine (Figure S3R-S). However, mass spectrometry showed an increase in the intensity of oxidized methionine peptides upon rotenone treatment (Figure S6F). We cloned a methionine-free version of tau, tau^Metless^, where every methionine is mutated to leucine. This construct was expressed at the same levels as WT tau but showed a large decrease in 25kD fragment formation (Figure 7E). Taken together, we conclude that direct oxidation of tau by ROS and ROS-induced changes in proteasomal processing contribute to the accumulation of the 25kD fragment.

### 25kD tau fragment changes tau aggregation

ROS accumulation and changes in proteasome activity are hallmarks of aging and often precede the detection of neurodegenerative disease-associated aggregates. We hypothesized that generation of the 25kD fragment, then, could alter the tau aggregation process. We purified tau 1-172 and 1-176 (Figure S2G). Next, we performed ThT assays using increasing concentrations of the 25kD fragment with 10µM 0N4R tau using heparin as an inducer (Figure 7F-G). The final ThT intensity as well as the lag time increased as a function of 25kD fragment concentration.

Aggregation of tau in the presence of BSA also increased the lag time, but did not increase the final ThT signal (Figure 7G). Negative stain electron microscopy revealed a straightening of the tau fibril quaternary structure as a function 25kD fragment concentration (Figure 7H-J). Therefore, the presence of the 25kD fragment can change tau fibril aggregation.

## Discussion

We established that *MAPT* V337M iPSC-derived neurons have higher levels of tau oligomers than *MAPT* WT iPSC-derived neurons. Based on this phenotype, we used CRISPR-based screens in iPSC-derived neurons to systematically uncover modifiers of tau oligomer and total tau levels. Our antibody-based screening methodology is widely applicable across many different cell types and antigens. We showcased the robustness of this methodology by performing seven additional small-scale CRISPRi screens across different antibodies and *MAPT* genetic backgrounds. For interactive datasets of all screens in this paper, please see CRISPRBrain.org^65^.

In comparison to other tau screens previously reported in the literature^72–76^, our data have broadly similar patterns of hit genes. We uncovered hit genes in expected pathways such as autophagy and the ubiquitin proteasome system (including *CUL5*), and unexpected pathways, including mitochondrial proteins, GPI-anchor biosynthesis, and UFMylation. A previous genome-wide screen for modifiers of tau levels performed in SHY5Y cells^74^ is generally in agreement with our screen and has several shared classes of genetic modifiers, including autophagy, mitochondrial function, and the UPS. Surprisingly, this screen identified *CUL5* as a negative modifier of tau levels, in contrast to our data (for a full comparison of these screens, see Figure S4B-D). Since CUL5 binds to at least 30 different substrate adaptors, it is not necessarily surprising that *CUL5* knockdown might have different phenotypes in different contexts.

We also screened our focused library in multiple secondary screens (Figure 3) and, as a collaborative project, in a seeding-based tau aggregation model in iPSC-derived neurons (Bravo et al.)^76^. The UFMylation pathway is also a strong modifier of tau seeding in that screen.

Surprisingly, in Bravo et al., nuclear-encoded mitochondrial genes are negative modifiers of tau seeding, while they are strong positive modifiers in this study. A full pair-wise comparison of the two screens is available in Figure S4E. Differential effects of genetic modifiers in different cell-based models of tauopathy highlight the fact that some cellular pathways may have opposing roles in promoting or inhibiting different forms of tau aggregation. Future studies will further investigate the biochemical and molecular mechanisms that underlie the activity of genetic modifiers of tau conformation to reconcile differences between model systems and to identify therapeutics for disease.

Here, we focused on the E3 ubiquitin ligase complex CRL5^SOCS4^. We find that CRL5^SOCS4^ ubiquitinates tau and controls tau levels specifically in the soma, but not the neurites. This suggests that CRL5^SOCS4^ mediates subcellular control of tau proteostasis. Synaptic impairment is thought to be due to tau accumulation in axons. However, NFTs in disease are canonically in the soma. Our data support a model where CRL5^SOCS4^ plays a housekeeping role for tau in the soma.

We find that the gene encoding the E3 ligase component, *CUL5,* is a novel factor that is correlated with vulnerability tauopathies as are genes encoding multiple CUL5 interactors, including *ARIH2* and *SOCS4*. *SOCS4* is expressed at higher levels in resilient neurons in the insula in a recent scRNA-seq study of FTD^12^. However, the molecular mechanism(s) by which *CUL5* affects neuronal vulnerability in AD remain to be identified. A broad distribution of *CUL5* (and *SOCS4*) expression is seen in different neuronal subtypes in the Seattle Alzheimer’s Disease Brain Cell Atlas^58^ suggesting that *CUL5* may modulate disease vulnerability via multiple mechanisms. For instance, it is possible that *CUL5* expression affects vulnerability via tau ubiquitination. But, considering *CUL5*’s known role in immune signaling, another possible hypothesis is that *CUL5* expression affects vulnerability via the neuro-immune axis. Deconvoluting cell-type specific CRL5 activity and *CUL5* expression will be necessary for understanding these mechanisms.

We also discovered a new connection between ROS, mitochondrial function, tau proteoforms, and tau aggregation; the strongest enriched KEGG pathway in our primary screen was oxidative phosphorylation. We found that acute inhibition of the ETC leads to ROS, which in turn leads to the generation and extracellular release of the NTA biomarker-positive 25kD tau fragment. This finding suggests that the NTA biomarker may report on neuronal oxidative stress in human patients. However, we lack direct evidence to prove this hypothesis. One tau fragment present in CSF, N-224, has been shown to be produced by calpain cleavage^67,77^. Our work reveals another mechanism by which N-terminally truncated tau could be generated in neurons: ROS-induced changes in proteasome activity coupled with direct oxidation of tau. The generation of this fragment directly links common observations in aging and neurodegeneration: decreases in mitochondrial function, proteasome activity, and changes to tau proteoforms. More work is needed to understand why tau oxidation results in changes to its proteolysis, including how tau oxidation changes tau conformation, and how tau degradation by the proteasome is altered. Beyond its utility as a biomarker, N-terminal tau fragments may also modulate tau aggregation, as suggested by our *in vitro* assays. Our data suggest multiple routes by which perturbation of the electron transport chain may contribute to tau oligomerization: (1) ROS generation (2) proteasome inhibition (3) tau proteolysis. Further work will be needed to provide the direct mechanisms both in model systems and in human disease.

Our work underscores the observation that cell types have diverse strategies for controlling tau conformation; how the proteostasis network interacts with tau can differ from cell type to cell type, and by subcellular compartment. Further work is needed to pinpoint how the molecular drivers that cause neurodegeneration modulate tau conformational changes that are evident in disease, and which of these drivers are suitable therapeutic targets.

## Limitations of the Study

This study has two main limitations: (1) reliance on anti-oligomer antibodies that are well characterized in disease, but lack high resolution structural information about the epitopes (with the exception of M204). (2) The use of iPSC-derived neurons.

A major limitation of this study is the lack of well-defined anti-oligomer antibody epitopes. The oligomeric structures many of these antibodies recognize are not well defined, and thus we were not able to perform a full structural or biochemical characterization of the oligomer species present in the *MAPT* V337M iPSC-derived neurons. Our biochemical data (dot blots; Figure 1A) show that all the antibodies had some signal in the *MAPT* WT and *MAPT* KD samples. This may be due to several factors, such as formation of oligomeric tau in *MAPT* WT cells, technical limitations in dot blot format, or cross-reactivity with monomeric tau. Therefore, our screen phenotypes could be interpreted as a combination of signal from oligomeric tau, monomeric tau, and non-specific signal. We overcame these limitations by validating the primary genome-wide screen hits with two more oligomer specific antibodies (Figure 3B-C).

The set of overlapping hits should be considered high confidence, and the counter screen for total tau levels highlights hits that likely affect monomeric tau levels as well. Our further mechanistic experiments focus on the mechanisms that are independent of the use of tau oligomer antibodies, and do not suffer from this limitation.

iPSC-derived neurons have known limitations when modelling neurodegenerative disease including, but not limited to: the exclusive expression of the fetal isoform of tau (0N3R), immature neuronal activity, a transcriptome more closely resembling immature neurons, and the inability to support spontaneous formation of tau fibrils. Furthermore, neuronal function and survival in disease is heavily modulated by other cell types in the brain such as glia, and in this study we use mono-cultured iPSC-derived neurons in a dish. However, comprehensive genetic modifier screens as well as mechanistic biochemical studies of tau have previously been performed mostly in cancer cells that do not recapitulate fundamental aspects of neuronal biology (including endogenous tau expression). In this way, the use of iPSC-derived neurons is a significant advance for modeling fundamental tau biology. Future work, including screens in preclinical mouse models of tauopathy using our recently establishing *in vivo* screening platform^78^, will be needed to translate our *in vitro* findings into useful strategies for understanding and treating the underlying molecular mechanisms that drive tauopathies.

## STAR ★ Methods

### KEY RESOURCES TABLE

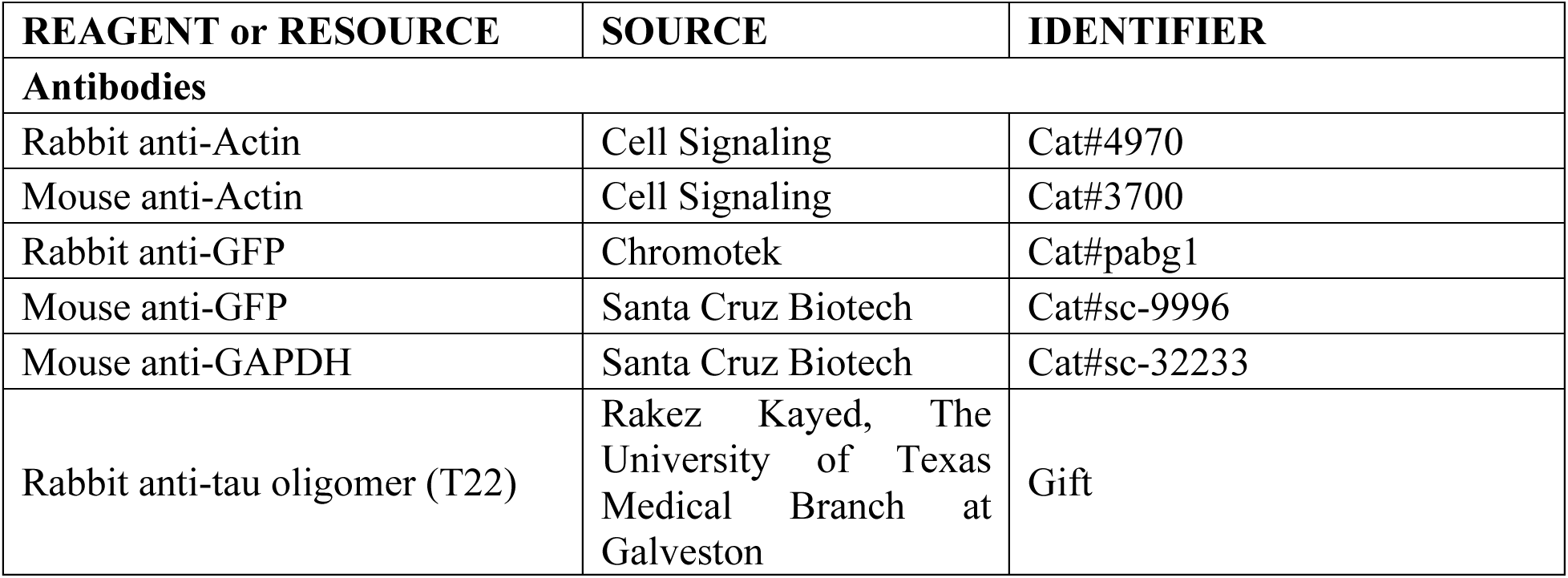
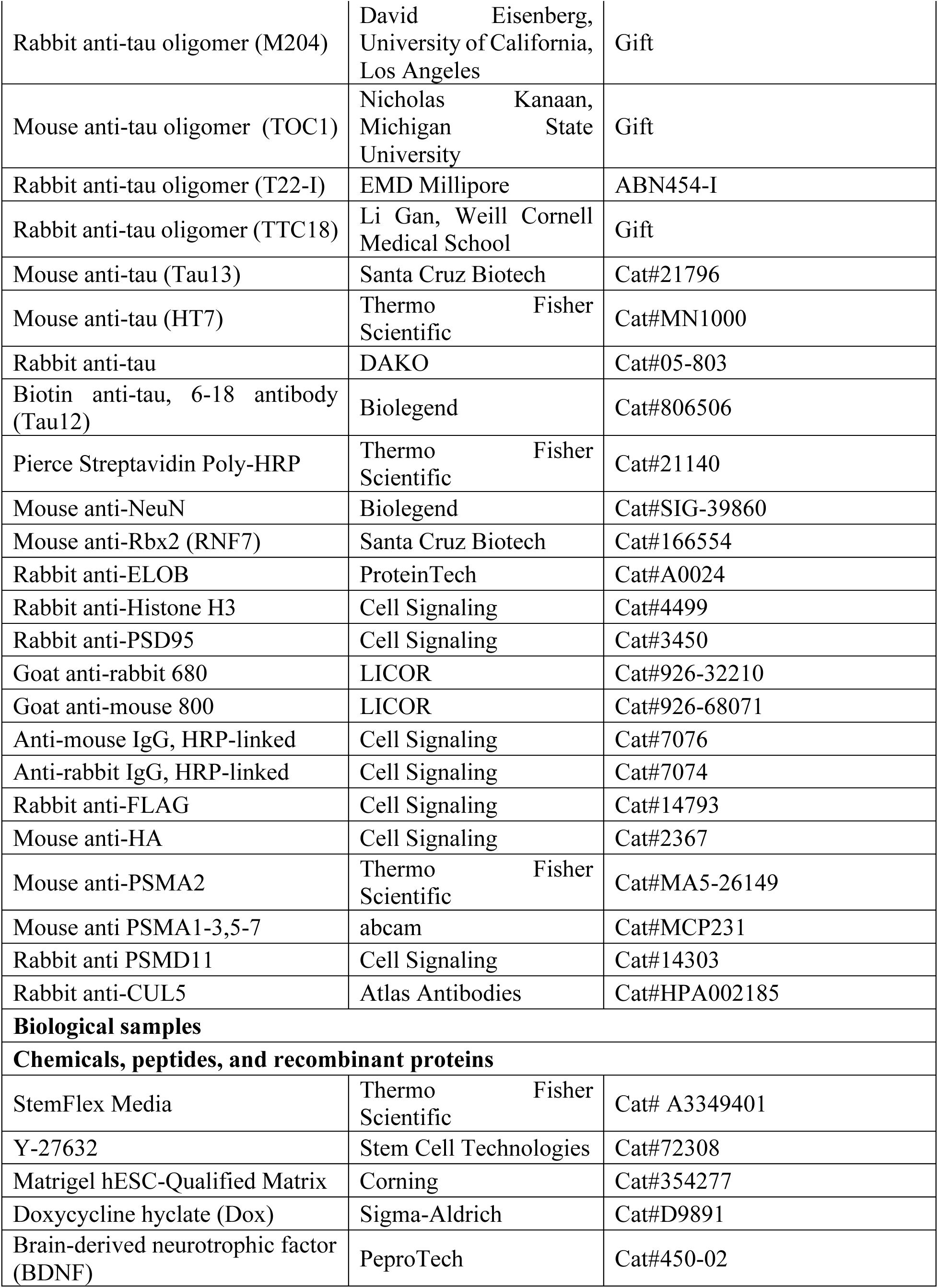
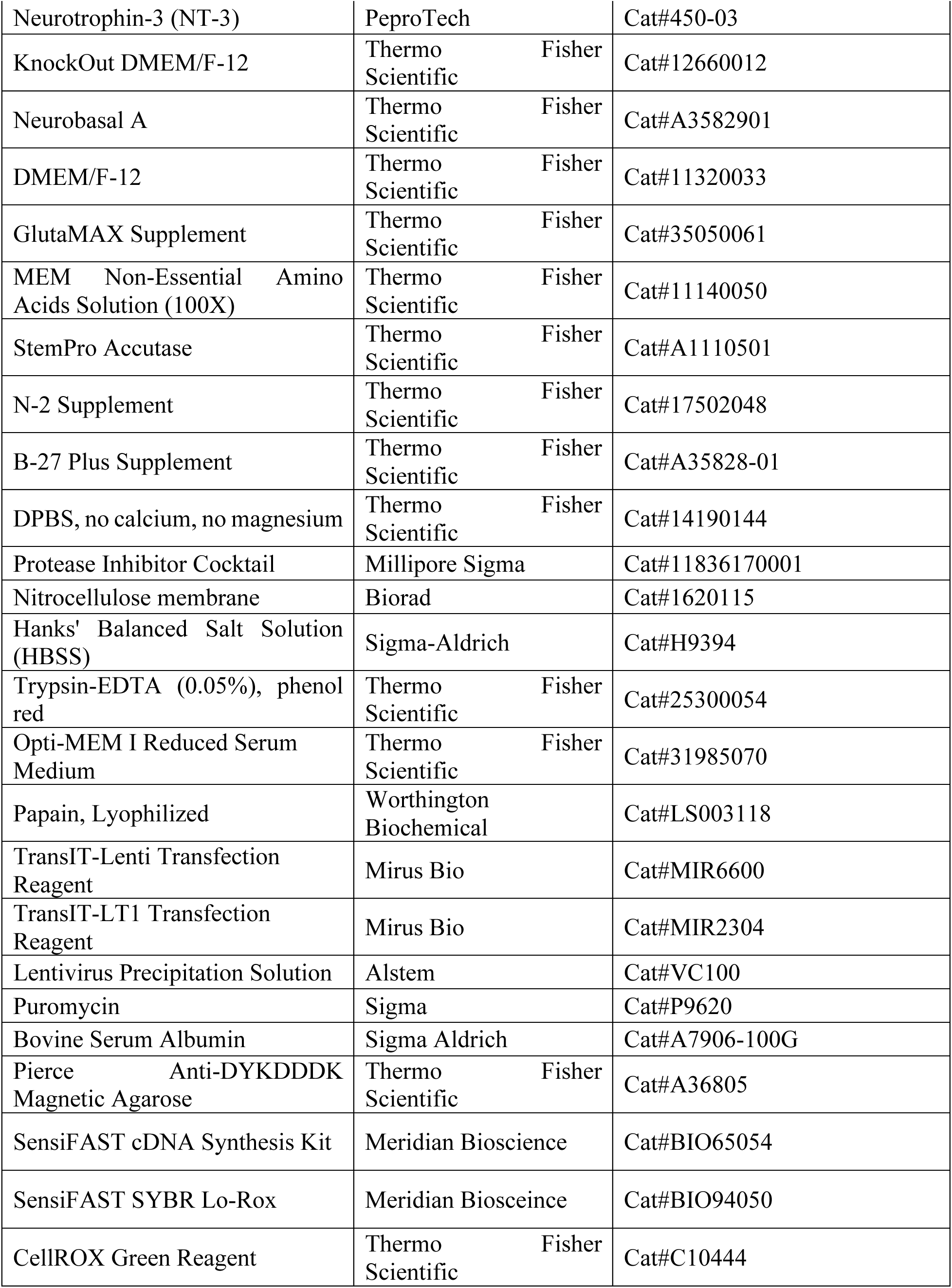
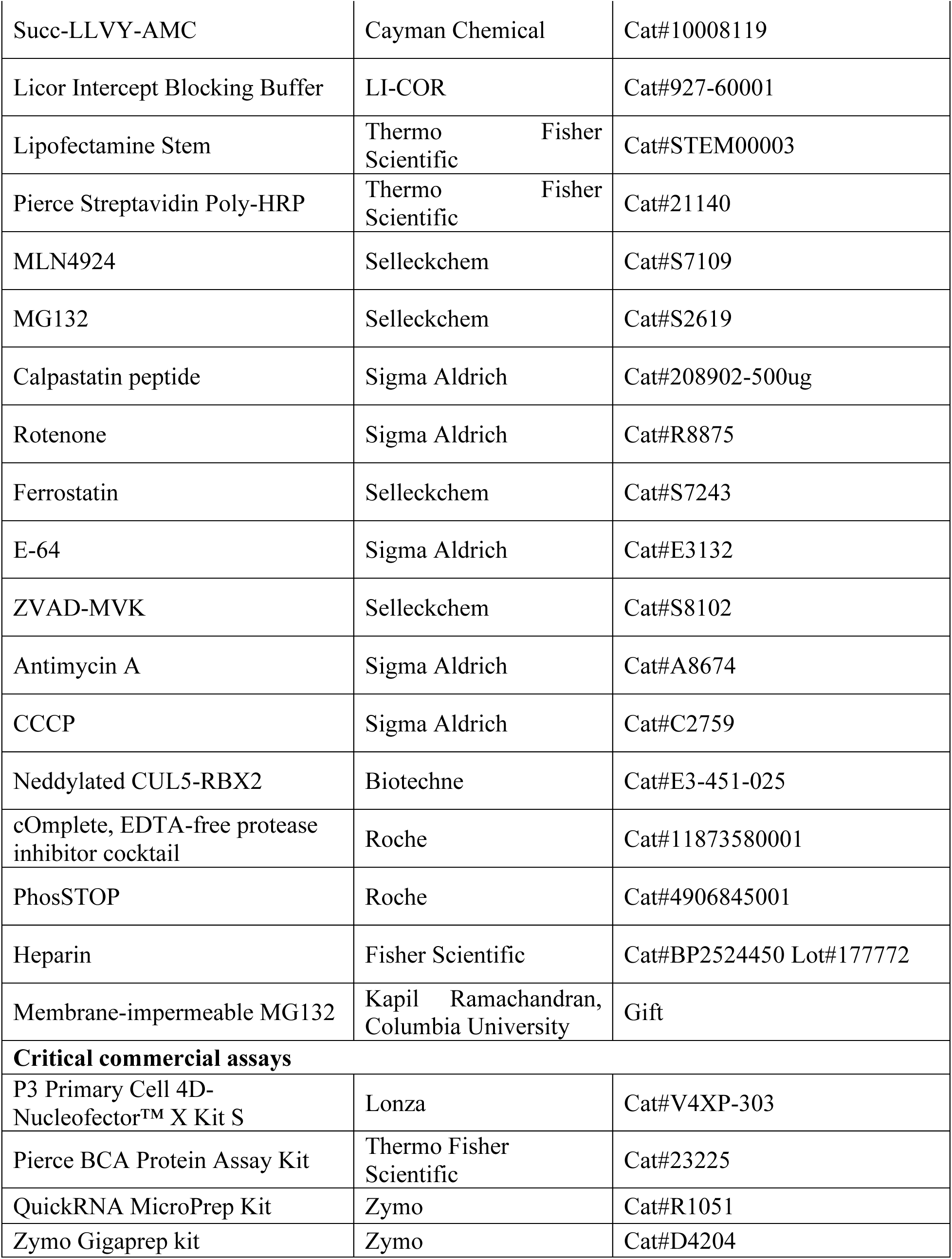
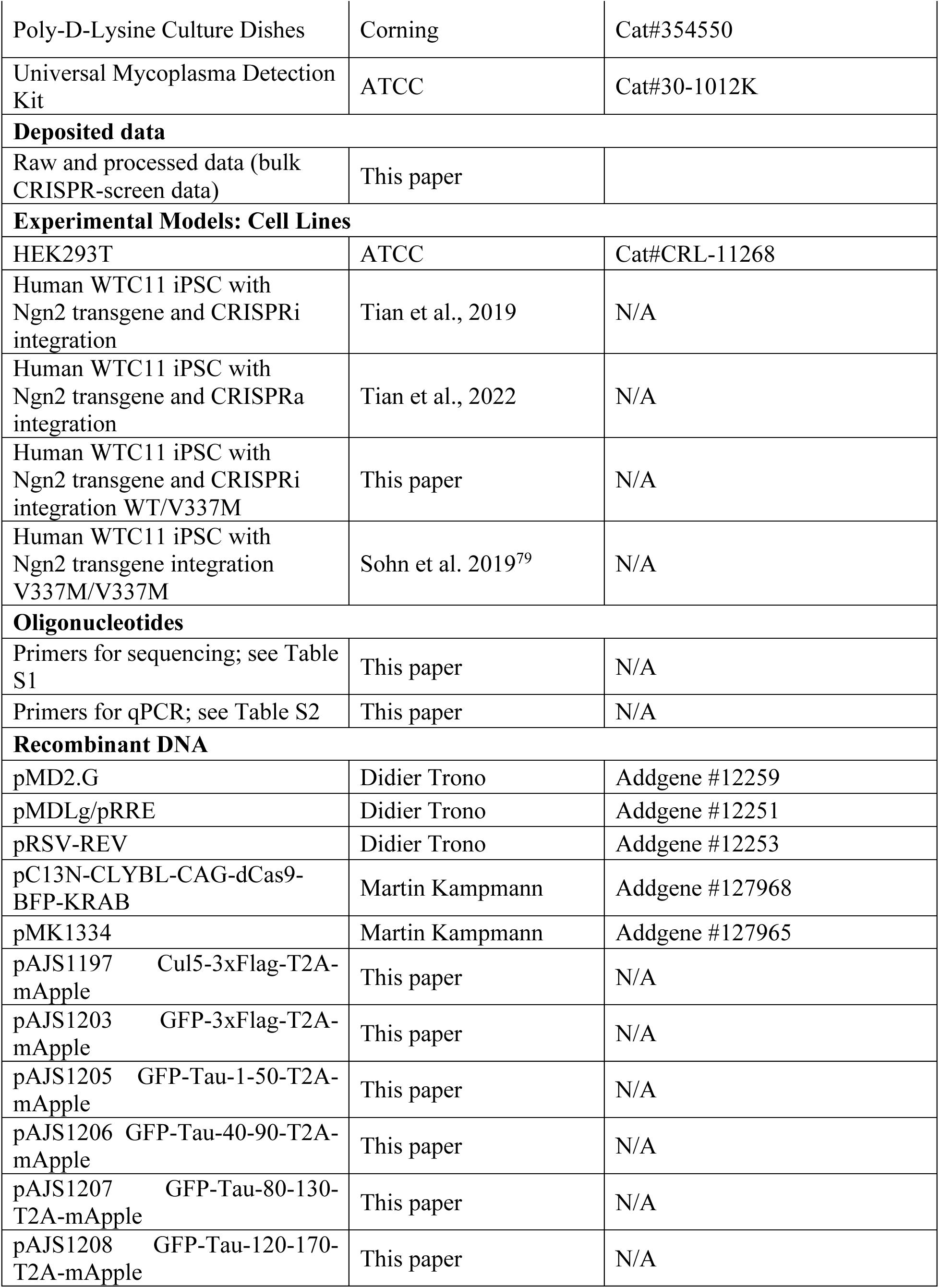
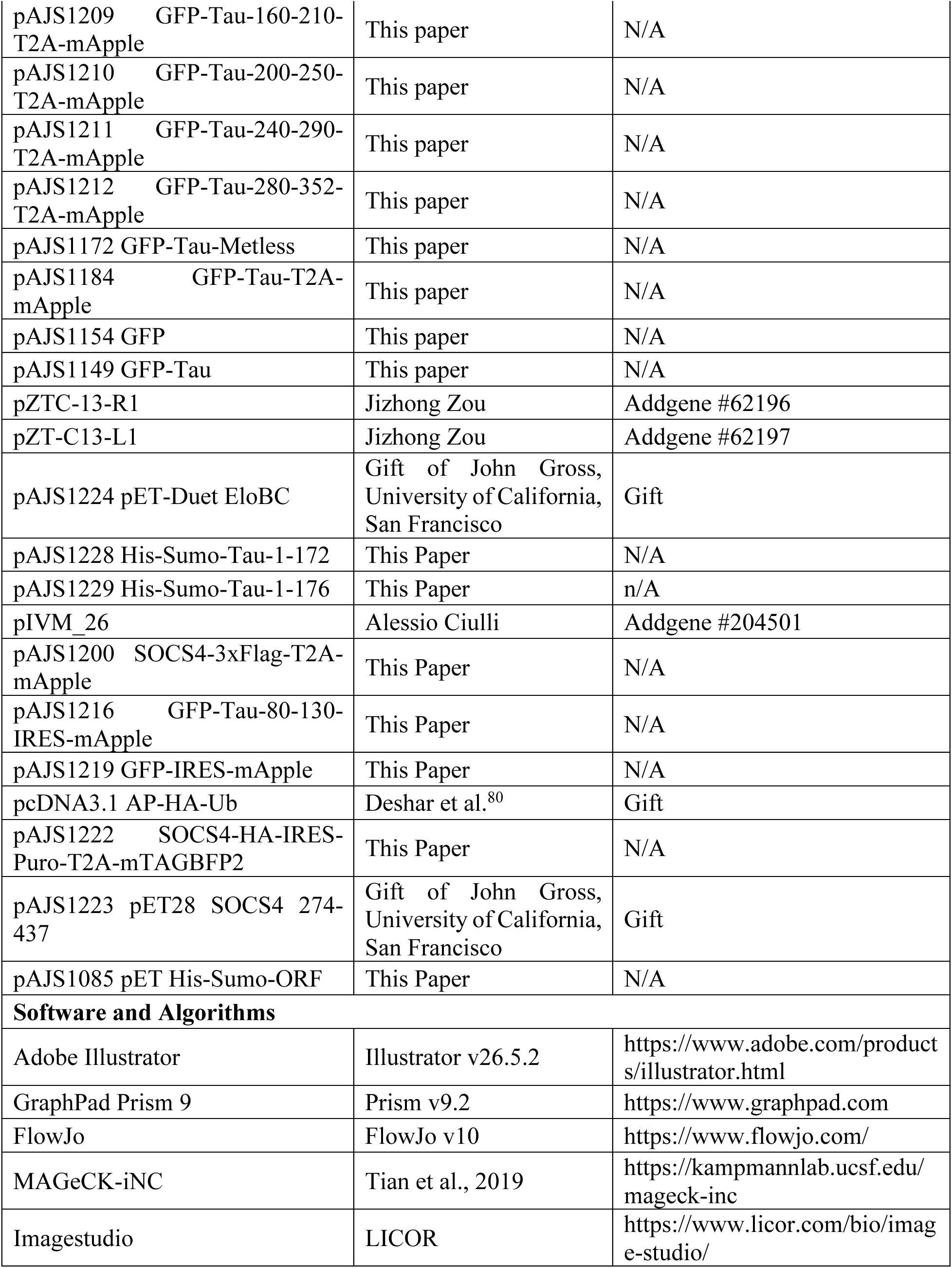

## RESOURCE AVAILABLILITY

### LEAD CONTACT AND MATERIALS AVAILABILITY

The cell lines generated in this study are available on request upon the completion of a Material Transfer Agreement (MTA). All plasmids generated in this study will be deposited on AddGene. Further information and requests for resources and reagents should be directed to and will be fulfilled by the Lead Contact, Martin Kampmann.

## DATA AND CODE AVAILABILITY

- CRISPR screening data will be deposited at CRISPR-brain (https://www.crisprbrain.org/) and will be publicly available as of the date of publication.
- CRISPR screening raw data will be made available upon request
- All original code is available on the Kampmann lab website (https://kampmannlab.ucsf.edu/resources)

## EXPERIMENTAL MODEL AND SUBJECT DETAILS

### Human iPSCs

Human iPSCs (in the male WTC11 background (Miyaoka et al., 2014)) were cultured in StemFlex Medium on BioLite Cell Culture Treated Dishes (Thermo Fisher Scientific; assorted Cat. No.) coated with Growth Factor Reduced, Phenol Red-Free, LDEV-Free Matrigel Basement Membrane Matrix (Corning; Cat. No. 356231) diluted 1:100 in Knockout DMEM (GIBCO/Thermo Fisher Scientific; Cat. No. 10829-018). Routine passaging was performed as described ^22^. Studies with human iPSCs at UCSF were approved by the The Human Gamete, Embryo and Stem Cell Research (GESCR) Committee. Informed consent was obtained from the human subjects when the WTC11 (Miyaokaet al., 2014) lines were originally derived.

### Human iPSC-derived neurons

Human iPSC-derived neurons were pre-differentiated and differentiated as described ^22^. Briefly, iPSCs were pre-differentiated in Matrigel-coated plates or dishes in N2 Pre-Differentiation Medium containing the following: KnockOut DMEM/F12 as the base, 1× MEM non-essential amino acids, 1× N2 Supplement (Gibco/Thermo Fisher Scientific, cat. no. 17502-048), 10 ng ml^−1^ of NT-3 (PeproTech, cat. no. 450-03), 10 ng ml^−1^ of BDNF (PeproTech, cat. no. 450-02), 1 μg ml^−1^ of mouse laminin (Thermo Fisher Scientific, cat. no. 23017-015), 10 nM ROCK inhibitor and 2 μg ml^−1^ of doxycycline to induce expression of mNGN2. After 3 d, on the day referred to hereafter as Day 0, pre-differentiated cells were re-plated into BioCoat poly-D-lysine-coated plates or dishes (Corning, assorted cat. no.) in regular neuronal medium, which we refer to as +AO neuronal medium, containing the following: half DMEM/F12 (Gibco/Thermo Fisher Scientific, cat. no. 11320-033) and half neurobasal-A (Gibco/Thermo Fisher Scientific, cat. no. 10888-022) as the base, 1× MEM non-essential amino acids, 0.5× GlutaMAX Supplement (Gibco/Thermo Fisher Scientific, cat. no. 35050-061), 0.5× N2 Supplement, 0.5× B27 Supplement (Gibco/Thermo Fisher Scientific, cat. no. 17504-044), 10 ng ml^−1^ of NT-3, 10 ng ml^−1^ of BDNF and 1 μg ml^−1^ of mouse laminin. Neuronal medium was half-replaced every week. Full protocols are available on protocols.io (dx.doi.org/10.17504/protocols.io.bcrjiv4n).

### HEK293T

HEK293Ts were cultured in DMEM supplemented with 10% FBS, 1% Pen/Strep, 1% Glutamine (DMEM complete). Cells were passaged by washing with DPBS, adding trypsin for 2 minutes at 37C, and quenching and resuspended in a 5-fold trypsin volume of DMEM complete. Cells were spun at 200xg for 5 minutes, counted, and plated at the desired density. Drug treatments were performed by changing to fresh media with the indicated drug for the indicated time.

## METHOD DETAILS

### Generation of V337M CRISPRi line

WTC11 WT/V337M iPSCs harboring a single-copy of doxycycline-inducible mouse NGN2 at the AAVS1 locus ^81^ were used as the parental iPSC line for further genetic engineering. iPSCs were transfected with pC13N-dCas9-BFP-KRAB and TALENS targeting the human CLYBL intragenic safe harbor locus (between exons 2 and 3) (pZT-C13-R1 and pZT-C13-L1, Addgene #62196, #62197) using DNA In-Stem (VitaScientific). After 14 days, BFP-positive iPSCs were isolated via FACS sorting, and individualized cells were plated in a serial dilution series to enable isolation of individual clones under direct visualization with an inverted microscope in a tissue culture hood via manual scraping. Clones with heterozygous integration of dCas9-BFP-KRAB (determined using PCR genotyping) were used for further testing. Full protocols are available on protocols.io: https://dx.doi.org/10.17504/protocols.io.8dahs2e.

### Flow cytometry

#### Fixed neurons

For CRISPR-screening, iPSC-differentiated neurons were washed with HBSS and then dissociated from the plate using Papain solution (20 U/mL papain in HBSS) at 37C for 10 minutes. Papain was quenched with 3x volume DMEM with 10% FBS and spun down 200xg for 10 minutes. Cells were then fixed with zinc fixation buffer (0.1M Tris-HCl, pH 6.5 @ 4C, 0.5% ZnCl2, 0.5% Zn Acetate, 0.05% CaCl2) overnight at 4C. When ready for staining, samples were washed three times in TBS by centrifugation at 200xg for 10 minutes and resuspended in permeabilization buffer (10% Normal Goat Serum, 10% 10x TBS, 3% BSA, 1% Glycine, 0.5% Tween-20) and blocked for 30 minutes. During blocking, cells were triturated into a single-cell suspension with progressively smaller pipette tips, from P1000 pipette tips to P20. After addition of primary antibodies, the antibody/cell slurry was incubated for two hours. Samples were then spun down at 200xg for 10 minutes and washed 3x with TBS by centrifugation. The cell pellet was then resuspended in permeabilization buffer with secondary antibodies and incubated at room temperature for one hour. Samples were then spun down at 200xg for 10 minutes and washed 3x with TBS by centrifugation. In the last wash, Hoechst at a concentration of 2 µM was added to the wash buffer for 10 minutes at room temperature. After the last wash, cells were resuspended in 1 mL of TBS and FACS sorted on a BD ARIA Fusion.

All other iPSC-differentiated neurons were prepared for flow cytometry in 96-well plate format as follows. iPSC-differentiated neurons were washed with TBS and then fixed with zinc fixation buffer at 4 °C overnight. When ready for staining, samples were washed three times in TBS and then 50uL of permeabilization buffer was added and incubated for 30 minutes to block. Primary antibody in permeabilization buffer was then added and samples incubated at 4 °C overnight.

Primary was removed and cells were washed 4x in TBS. Secondary antibodies were added in permeabilization buffer and incubated for one hour at room temperature. Cells were washed 4x in TBS with Hoechst in the second to last wash. After the last wash, cells were triturated by pipetting up and down with a P200 tip 10 times and then a P20 tip 10 times and analyzed with a BD Celesta cell analyzer and the data were processed using FlowJo v10 software.

### Live neurons

iPSC-differentiated neurons were washed with HBSS and dissociated from the plate using Papain solution (20 U/mL papain in HBSS) at 37 °C for 10 minutes. Papain was quenched with 3x volume DMEM with 10% FBS and spun down 200xg for 10 minutes. Cells were then resuspended in HBSS and analyzed with a BD Celesta analyzer the data were processed using FlowJo v10 software. For cells analyzed with CellRox (ThermoFisher Scientific), CellRox diluted to 5 µM in differentiation media was added 1:1 with the extant well media volume and incubated for 30 minutes at 37 °C. Cells were washed with HBSS three times and dissociated and flowed as described above.

### HEK Cells

HEK cells were washed with DPBS and dissociated with 0.25% Trypsin EDTA for 2 minutes at 37 °C. Trypsin was quenched by addition of a five-fold volume of DMEM Complete and spun down for 5 minutes at 200x*g*. Cells were resuspended in DPBS +0.5% FBS, filtered through a blue-capped FACS tube (35 µm strainer) and subjected to flow cytometry.

### Dot Blotting

iPSC-derived neurons were grown as described. Cells were washed 3x with DPBS and scraped down into 1mL of DPBS with protease and phosphatase inhibitors. Cells were homogenized in a Dounce homogenizer for 30 strokes on ice and spun down at 21,000x*g* for 10 minutes at 4 °C. the DPBS soluble supernatant was removed and the pellet was then resuspended in DPBS+0.1% Triton X-100, triturated and spun again at 21,000xg for 10 minutes at 4 °C. The 0.1% Triton X- 100 soluble supernatant was removed and the pellet was then resuspended in DPBS+0.5% SDS, triturated and spun again at 21,000xg for 10 minutes at room temperature. Supernatants were then blotted onto nitrocellulose membranes and allowed to dry for one hour at room temperature. Blotting proceeded as described for western blotting above, except using 5% non-fat dry milk as a block.

### Lentivirus generation

#### sgRNAs

Lentivirus was generated as described ^22^. Briefly, HEK293T were plated to achieve 80-95% confluence 24 hours after plating. For a 6-well plate, 1ug of transfer plasmid, and 1ug of third generation packaging mix, were diluted into 200 µL OPTIMEM and 12 µL of TRANSIT-LENTI was added. Transfection mix was incubated at room temperature for 10 minutes and then added to HEKs. After 48 hours, supernatant was transferred to a syringe and filtered through a 0.45 µm PVDF filter into a conical tube. ¼ volume of Lentivirus Precipitation Solution was added to the filtrate and stored at 4 °C for 24 hours. Lentivirus-containing supernatant was then centrifuged for 30 min at 1500xg at 4 °C, supernatant aspirated, and then spun again at 4 °C for 5 mins at 1500xg. Supernatant was aspirated and the virus-containing pellet was resuspended in 200 µL DPBS. For screening, each library was prepared from a 15cm plate of HEK293Ts, scaled appropriately. For infection, virus was added at the same time as iPSC passaging. After 48 hours, cells were passaged, analyzed for marker positivity by flow cytometry and selected with 1 µg/mL puromycin until >95% marker positive, for two passages. Cells were allowed to recover for one passage before pre-differentiation. The full protocol is available on protocols.io (https://dx.doi.org/10.17504/protocols.io.8dfhs3n)

### Over-expression constructs

Lentivirus was generated as described ^22^. Briefly, HEK293T were plated to achieve 80-95% confluence 24 hours after plating. For a 6-well plate, 1 µg of transfer plasmid, and 1 µg of third-generation packaging mix, were diluted into 200uL OPTIMEM and 12 µL of TRANSIT-LENTI was added. Transfection mix was incubated at room temperature for 10 minutes and then added to HEK293T cells. After 48 hours, supernatant was transferred to a syringe and filtered through a 0.45 µm PVDF filter into a conical tube. ¼ volume of Lentivirus Precipitation Solution was added to the filtrate and stored at 4 °C for 24 hours. Lentivirus-containing supernatant was then centrifuged for 30 min at 1500xg at 4 °C, supernatant aspirated, and then spun again at 4 °C for 5 mins at 1500xg. Supernatant was aspirated and the virus-containing pellet was resuspended in 200n µL DPBS. Cells were expanded to T75 flasks and when 80% confluent sorted for the appropriate marker (mApple or GFP).

### CRISPRi screening

For each genome-wide sub-library and each secondary screen, 45 million iPSCs in 3x T175s were infected with lentivirus as above at an MOI of ∼0.3 and selected. Cells were then differentiated as above and plated on 3x 15-cm PDL-coated dishes at a density of 15 million cells per plate. Cells were then matured for two weeks and prepared for FACS sorting as above, staining for NeuN and the tau-specific antibodies as indicated. For the tau-specific antibodies with mouse host, NeuN staining was not performed. Cells were collected into 1mL of 30% BSA in a FACS tube. After sorting, cells were pelleted at 200xg for 20 minutes, the supernatant was removed and the pellet was frozen at -20. Genomic DNA was extracted with the NucleoSpin Blood L kit. sgRNA cassettes were amplified pooled and sequenced as described ^22^. Sequencing was analyzed as described for each sub-library ^22^. Screening data is available in Supplemental Tables 2 and 3.

### Cloning of secondary screen library

A pool of sgRNA-containing oligonucleotides were synthesized by Agilent Technologies and cloned into our optimized sgRNA expression vector as previously described^82^.

### Mechanical fractionation of iPSC-derived neurons

Neurons were mechanically fractionated as described^53^. Briefly, neurons plated on day 0 on PDL-coated microporous transwell membranes, 1 µm pore diameter (Corning 353102) in PDL-coated 6-well cell culture plates (Corning 353502) and fed as described above. On day 14, transwell inserts were removed and the top of the membrane was scraped into lysis buffer. The bottom of the membrane was then cut out of the transwell plate and scraped into lysis buffer. Downstream processing was performed as below.

### Western Blotting

iPSC-derived neurons were cultured as described above. Neurons were washed 3x with ice-cold DPBS and then ice-cold RIPA with protease and phosphatase inhibitors was added to cells (50 µL for a 24-well plate). Lysates were incubated on ice for 2 minutes and then scraped down.

Lysates were either flash frozen on liquid nitrogen or directly centrifuged at 21000xg for 10 minutes at 4 °C. The supernatants were then collected, and concentrations assessed using a BCA assay (Thermo). 10 µg protein were loaded onto a 4-12% Bis-Tris polyacrylamide gel (Thermo). Nitrocellulose membranes were used to transfer the protein in a BioRad Transblot for 11 minutes at 25 V, 25 A. Membranes were then blocked for 1 hour with Licor Odyssey block and primary was added in Licor Odyssey block overnight at 4 °C. Blots were then washed 4x 5 minutes with TBST and secondary antibodies were added in Licor Odyssey block for 1 hour at room temperature. Blots were washed 4x 5 minutes with TBST and imaged on a Licor. Immunoblots were quantified by intensity using ImageStudio (Licor).

### Immunoprecipitations

#### Flag IPs

iPSC-derived neurons were cultured as described above. Neurons were washed 3x with ice-cold DPBS and then lysed in FLAG-lysis buffer (20 mM HEPES NaOH pH 7.4, 150 mM NaCl, 0.2% NP40 with protease and phosphatase inhibitors and 2 µM 1,10-phenathroline). Lysates were freeze/thawed on liquid nitrogen 7 times and then centrifuged at 21000xg for 30 minutes at 4C. The supernatants were then collected and concentrations assessed using a BCA assay. 5 mg of lysate was loaded onto 25 µL FLAG dynabeads (thermo) that had been washed 3x with FLAG-lysis buffer. IP was performed with rotation at 4 °C for 2 hours. Beads were then washed 3x with FLAG-lysis buffer and then 2x with TBS. FLAG-peptide elution was performed with 1 mg/mL FLAG-peptide in TBS overnight at 4C. Acid elution with 100 mM glycine pH 2.0 was performed for 10 min and the supernatant was quenched in 1/10^th^ volume of 1M Tris pH 9.0. High-temperature elutions were performed using 2x LDS with 20 mM DTT for 70 °C for 5 minutes.

### GFP IPs

iPSC-derived neurons were cultured as described above. Neurons were washed 3x with ice-cold DPBS and then lysed in GFP-lysis buffer supplemented with protease and phosphatase inhibitors (ProteinTech). Lysates were freeze/thawed on liquid nitrogen 7 times and then centrifuged at 21000xg for 30 minutes at 4 °C. The supernatants were then collected and diluted using GFP dilution buffer supplemented with protease and phosphatase inhibitors (ProteinTech).

Concentrations were assessed using a BCA assay. 100 mg of lysate was loaded onto 100 µL GFP-trap beads (ProteinTech) that had been washed 3x with Dilution buffer supplemented with protease and phosphatase inhibitors. IP was performed with rotation at 4 °C for 1 hours. Beads were then washed 3x with GFP-wash buffer and then 2x with TBS. Acid elution with 100 mM glycine pH 2.0 was performed for 10 min and the supernatant was quenched in 1/10^th^ volume of 1M Tris pH 9.0. Eluates were then loaded on a gel and excised for mass spectrometry (below).

HEK cells were cultured as described above. Cells were washed 3x with DPBS and lysed in RIPA plus 10% glycerol, 50mM iodoacetamide (a de-ubiquitinase inhibitor), 2.5 mM MgCl_2_, 125U Benzonase, and protease and phosphatase inhibitors for 30 minutes on ice. Lysates were then spun down at 21,000x*g* for 10 minutes at 4 °C. Supernatants and pellets were separated. Supernatant concentrations were calculated by BCA and normalized. Inputs were then loaded onto 25 µL of GFP-trap beads that were pre-equilibrated in lysis buffer. IP was performed with rotation at 4 °C for 2 hours. Beads were then washed 2x with GFP wash buffer (RIPA +10% glycerol) then high salt GFP wash buffer (RIPA with 0.5 M NaCl +10% glycerol), and then once more in GFP wash buffer. Beads were then resuspended in 75 µL of 2x SDS-PAGE loading dye and boiled for 5 minutes at 95 °C. Eluates were then separated from the beads using the magnetic and run on Western blots as above.

### HEK Cell Transfections for immunoprecipitations

Cells were seeded the day before transfection at 1x10^6^ per well in a six well plate. The next day, for each well, 2000 ng of GFP or GFP-tau containing construct, 2000 ng of SOCS4-Flag, and 500 ng of HA-Ubq were mixed in 200 µL OPTIMEM. To this, 7.5 µL of Mirus LT1 was added and incubated for 10 minutes. Cells were refed with fresh DMEM Complete during this time.

After ten minutes, transfection mix was added to cells. Twenty-four hours later, media was exchanged with or without drug treatment and harvested for IP after the indicated drug treatment time.

### In-gel proteasome assay

Proteasome activity from neurons was measured as described (<u>10.1016/j.xpro.2021.100526</u>). Briefly, iPSC-derived neurons were washed 3x with ice cold DPBS, and then scraped into TSDG lysis buffer (10 mM Tris, 1.1 mM MgCl2, 10 mM NaCl, 1 mM NaN3, 1 mM DTT, 2 mM ATP, 10% glycerol) and lysed by freeze/thaw on liquid nitrogen 7 times. Samples were then spun down 10 minutes at 21000xg, assayed by BCA, and 50 µg supernatant was mixed with 5x native gel loading buffer (0.05% bromophenol blue, 43.5% glycerol, 250 mM Tris pH 7.5) to a concentration of 1x loading buffer. Samples were loaded on a 3-8% Tris Acetate gel (Thermo) was run in native gel running buffer (1x TBE, 413 µM ATP, 2 mM MgCl2, 0.5 mM DTT) for 3 hours at 170 V. Gels were then incubator for 30 min at 37 °C in reaction buffer (50 mM Tris pH 7.5, 10 mM MgCl2, 1 mM ATP, 1 mM DTT, 48 µM Suc-LLVY-AMC) and imaged on a Gel Doc EZ (Bio-rad).

### Mass spectrometry

#### Identification of 25kD Fragment

Gel bands were then excised and the proteins making up each band were reduced, alkylated, and digested with trypsin all within the gel. The resulting peptides were then extracted from the gel slice into solution and analyzed using liquid chromatography tandem mass spectrometry (LC-MS/MS) with data-dependent acquisition. Preparation in solution was used because alternate proteases, such as GluC, cannot penetrate the gel matrix making gel separation and preparation infeasible. After acquiring and analyzing our LC-MS/MS data, we identified three peptides that were semi-specific for the GluC with tryptic N termini, ending at 172 and 176 in the fetal MAPT sequence.

### Identification of ubiquitination sites

*In vitro* ubiquitin reactions were diluted 1:16 with ArgC protease activation buffer (50mM Tris HCl 7.5, 5 mM CaCl_2_, 2mM EDTA, 5 mM DTT) before ArgC protease was added at a 1:25 protease:protein ratio. Samples were digested at 37°C for 3 hours with 400 rpm shaking. After digestion samples were desalted using NEST UltraMicroSpin columns per manufacturer’s instructions. After elution from desalting columns, samples were frozen and dried in a Speed- Vac centrifugal concentrator. Dried samples were resuspended in 50 µl of 0.1% formic acid before filtering through 0.45 µm filter and injection onto the Thermo Scientific Vanquish Neo HPLC platform on-line with a Thermo Exploris 480 Orbitrap Mass Spectrometer.

Peptides were separated using a Bruker 15 cm long 150 µm ID PepSep column packed with 1.5 µm BEH particles, over a 45 min gradient with mobile phase A composed of 0.1% formic acid in water and mobile phase B composed of 0.1% formic acid in 80% acetonitrile. The chromatographic gradient ran at a 600 nl/min flow rate throughout. The gradient started at 4% B before increasing to 28% B over 30 minutes, followed by an increase to 45% B over 5 minutes, and finally finishing with a wash of 95% B for 9 minutes. Full scans were collected at a resolution of 120,000 with a normalized AGC target of 100% and the maximum injection time set to “Auto”. Data dependent scans were collected at a resolution of 15,000 with a normalized AGC target of 200% and maximum injection time set to “Auto”. Precursors were selected for sequencing based on an allowed charge state of 2-6, and a dynamic exclusion after two sequencing events for 20 seconds of precursors within 10 ppm. The total cycle time of the full scan and all dependent (MS2) scans was 1 second.

Data was searched using MSFragger within the Fragpipe environment using default DDA settings searched against a full human proteome with the ubiquitin remnant as a variable modification and no carbamidomethylation of cysteine.

### ELISA

Capture antibody (either HT7 or DAKO, see Key Resources) were resuspended to a final concentration of 28ug/mL in Carbonate Buffer (0.2M carbonate/bicarbonate pH 9.4) and 100µL of this solution was used to coat each well for 1 hour at 25°C. After coating, wells were washed five times with 300µL/well of TBST (TBS+0.1% Tween-20). Wells were then blocked with 200µL of StartingBlock (ThermoFisher Scientific cat#37542) for 1 hour at 25°C. Wells were then emptied and for each well, 100uL of neuron conditioned media from one-well of a 96-well plate was added or purified tau standards. Samples were then incubated overnight at 4°C on a rocker. The next day, wells were washed five times with 300µL/well of TBST. 100µL of detection antibody (Biotinylated Tau12, see Key Resources), was added at 5µg/mL in StartingBlock for 2 hours at 25°C. Wells were then washed five times with 300µL/well of TBST. 100uL of Pierce Streptavidin Poly-HRP diluted 1:10,000 in StartingBlock was added for 1 hour at 25°C. Wells were then washed five times with 300µL/well of TBST. Supersignal ELISA Femto Substrate (ThermoFisher Cat#37075) was added to each well according to the manufacturer’s instructions. Luminescence was read on a Spectramax M5 microplate reader (Molecular Devices) with a 500ms integration time.

### qPCR

RNA was extracted using the Zymo Quick-RNA miniprep kit and cDNA was synthesized with the SensiFAST cDNA synthesis kit. Samples were prepared for qpCR in technical triplicates using SensiFAST SYBR Lo-ROX 2x Mastermix. qPCR was performed on an Applied Biosystems Quantstudio 6 Pro Real-Time PCR System using Quantstudio Real Time PCR Software following Fast 2-Step protocol: (1) 95 °C for 20 s; (2) 95 °C for 5 s (denaturation); (3) 60 °C for 20 s (annealing/extension); (4) repeat steps 2 and 3 for a total of 40 cycles; (5) 95 °C for 1 s; (6) ramp 1.92 °C s^−1^ from 60 °C to 95 °C to establish melting curve. Expression fold changes were calculated using the ΔΔCt method, normalizing to housekeeping gene *GAPDH*. Primer sequences are provided in Supplementary Table 1. Quantification of qPCR samples are shown in Figure S5.

### Drug treatments

At the two-week feeding, 50% of the media volume was removed and drugs diluted in media were added in order to obtain the desired drug concentration when adding media to reach the final media volume.

### Protein purification

UbE1, CDC34, UbE2L3, Neddylated Cul5/Rbx2, NEDD8 E1, NEDD8 E2, NEDD8, ARIH2, were all prepared and purified as previously described^83^.

### SOCS4

Full-length SOCS4 was amplified from iPSC cDNA and subcloned into pAJS1223 using NdeI and XhoI sites. SOCS4 ΔIDR (pAJS1223) was a gift (see start methods table). SOCS4 and EloBC constructs were transformed into BL21 Rosetta2 cells. 10 mL of an overnight starter culture grown in LB was used to inoculate 1L of TB and grown to an OD of 0.6 at 37 °C. Cultures were then moved to 18 °C and induced with 0.5mM IPTG. Cultures were grown for 12- 16 hours at 18 °C before spinning down at 4000x*g* for 10 minutes. All purification steps were carried out at 4 °C. Pellets were resuspended in SOCS4 lysis buffer (50 mM HEPES NaOH pH 7.5, 50 mM imidazole pH 8.0, 150 mM NaCl, 1 mM TCEP), lysed for 6 cycles of 30 seconds on/30 seconds off by a tip sonicator at 30% power. Lysates were spun down for 30 minutes at 30,000xg and loaded onto a 5-mL FF nickel column and eluted with a 50 mM-to-500 mM imidazole gradient. Fractions were run on a gel, pooled and dialyzed into lysis buffer overnight in the presence of 1.0 mg TEV protease. This sample was run over a second nickel column and the flowthrough was collected, dialyzed into lysis buffer, and run on a Q column eluted with a gradient from 150 to 500 mM NaCl. Fractions containing ternary complex were pooled, concentrated and run on an S200 (or S75 for SOCS4ΔIDR) pre-equilibrated in 25 mM HEPES pH 7.5, 250 mM NaCl, 10% glycerol, 2 mM DTT. Fractions containing ternary complex were concentrated and pooled.

### Tau 0N3R 1-172 and 1-176

Tau fragments 1-172 and 1-176 were subcloned into pAJS1085 (a His-Sumo-ORF) bacterial expression vector derived from pET28a using BsaI and BamHI restriction sites. These plasmids (pAJS1228 and pAJS1229) were transformed into BL21 Rosetta2 cells. 10 mL of an overnight starter culture grown in LB was used to inoculate 1L of LB and grown to an OD of 0.6 at 37 °C. Cells were induced for 4 hours at 37 °C with 1 mM IPTG and spun down at 4000x*g* for 10 minutes. Pellets were resuspended in Tau lysis buffer (50 mM HEPES NaOH pH 7.5, 50 mM imidazole pH 8.0, 150 mM NaCl, 2 mM TCEP), lysed for 6 cycles of 30 seconds on/30 seconds off by a tip sonicator at 30% power. Lysates were spun down for 30 minutes at 30,000x*g* and loaded onto a 5-mL FF nickel column and eluted with a 50 mM-to-500 mM imidazole gradient. Fractions were run on a gel, pooled and dialyzed into lysis buffer overnight in the presence of 0.25 mg Ulp1 protease. This sample was run over a second nickel column and the flowthrough was collected, concentrated, and run on a S75 column equilibrated in DPBS. Fractions were pooled, concentrated and flash-frozen for future use.

### Tau dGAE

Expression of tau was essentially performed as previous described^84^. Tau dGAE was transformed into *E. coli* BL21 (DE3) gold cells. A single colony was inoculated into 50 mL Terrific Broth (TB) with 50 mg/mL kanamycin as a starter culture and grown overnight at 37°C. The starter culture was then diluted into 4 L of TB and grown to an OD600 of 0.6 before inducing with 1mM IPTG at 37°C for 4 hours. The cells were harvested by centrifugation (4000x g for 20 min at 4°C), and flash frozen. Flash frozen pellets were resuspended in washing buffer (WB: 50 mM MES at pH 6.0; 10 mM EDTA; 10 mM DTT; 20 mM NaCl, supplemented with 0.1 mM PMSF and cOmplete EDTA-free protease cocktail inhibitors, at 10 ml/g of pellet). Cell lysis was performed using sonication (at 40% amplitude for 10 min, 5 s on/10 s off). Lysed cells were boiled at 80°C for 20 minutes, centrifuged at 20,000x g for 20 minutes at 4°C, and filtered through 0.45 μm cut-off filters and loaded onto a HiTrap SP FF 5 mL column (Cytiva Life Sciences) for cation exchange. The column was washed with 10 column volumes of WB and eluted using a gradient of WB containing 0.01-1 M NaCl. Fractions of 2 mL were collected and analyzed by SDS-PAGE. Proteins containing fractions were pooled and precipitated using 0.3 g/mL ammonium sulphate and left on a rocker for 30 min at 4°C. Precipitated protein was then centrifuged at 20,000xg for 30 min at 4°C and resuspended in 3 mL Dulbecco’s phosphate-buffered saline (DPBS) with no calcium or magnesium (Gibco) and desalted into 4 mL of 20 mM phosphate buffer at pH 7.2. The desalted protein was then concentrated to 14 mg/mL using molecular weight concentrators with a cut-off filter of 5 kDa. Purified protein samples were flash frozen in 100 uL aliquots for future use.

#### *In vitro* ubiquitination assays

*In vitro* ubiquitination was performed essentially as previously described^83^. Briefly, Tau constructs and SOCS4 were mixed with neddylated Cul5/Rbx2 at a 1:1 molar ratio 30 minutes prior to combining the substrates-E3 mixture, and charging reactions. Reactions were quenched with protein loading buffer containing SDS and betamercaptoethanol.

### ThT Assays

ThT assays were performed as described previously with heparin^85^, with the addition of the indicated concentrations of tau fragment constructs.

### Analysis of SEA-AD dataset

We utilized the SEA-AD single nucleus RNA sequencing dataset^58^ to determine (1) how *CUL5* expression of CUL5 and genes encoding proteins known to form CRL5 complexes changed across neocortical cell types in AD and (2) whether they were differentially changed in vulnerable (those that decrease in their relative abundance in AD) versus unaffected cell types. The datasets’ 84 donors span the spectrum of AD pathology and each is assigned a continuous disease pseudo-progression score (CPS) based on their extent of pathology measured by immunohistochemistry (against amyloid and hyperphosphorylated tau). For each cell type, we obtained the mean expression of each gene as well as effect sizes for how each gene changed along CPS (estimated with NEBULA^86^). We then correlated these values to effect sizes for how the cell types changed in their relative abundance along CPS (estimated with scCODA^87^). We then obtained a Pearson correlation coefficients between these values (and associated p-values) using the pearsonr function from scipy stats (version 1.12.0). P-values were adjusted for multiple hypothesis testing with the false_discovery_control function, also from scipy stats. The regplot function from seaborn (0.11.2) was used to plot a linear regression, which also computed 95% confidence interavls across 1,000 bootstraps.

### Pseudotime analysis with SlingShot

As in Rexach et al.^12^, we applied Slingshot 2.2.0^88^ to layer 2/3 IT excitatory neuron from the insular cortex for a total of 12218 cells across 37 samples. A UMAP-based dimensional reduction was calculated on the first 30 principal components of the scaled expression data. UMAP data and subcluster labels were supplied as input to Slingshot 2.2.0, generating an imputed lineage and pseudotime values per cell. Pseudotime was correlated with gene expression through Pearson correlation. P-values were adjusted with FDR correction across the number of genes.

### Negative Stain EM

Samples were taken from the endpoints of reaction conditions used for ThT assays. Samples were negatively stained with 0.75% uranyl formate (pH 5.5-6.0) on thin carbon-layered 400- mesh copper grids (Pelco) as described^89^. Negative stain EM micrographs were collected on a Talos L120C transmission electron microscope (FEI) equipped with a LaB6 filament operated at 120 keV. Images were recorded at a magnification of 54901x on a Ceta 16M detector with a 4k x 4k CMOS sensor with a 2.55 Å per pixel spacing.

## QUANTIFICATION AND STATISTICAL ANALYSIS

For statistical analysis, we used GraphPad Prism 9.5.1. Data are shown as mean±SD, except for flow cytometry data, which is shown as the median±SD. For two sample comparisons, un unpaired two-tailed Student’s t-test was used. For three sample comparison, a two-way ANOVA was used. P-values are shown above compared samples, n.s. denotes not significant.

### CRISPR-screen analysis

#### Primary screen analysis

CRISPR screens were analyzed using MAGeCK-iNC as previously described^22^. Briefly, raw sequencing reads were cropped and aligned using custom scripts that are already publicly available (https://kampmannlab.ucsf.edu/resources). Raw phenotype scores and p-values were calculated for target genes and negative control genes using a Mann-Whitney U-test. Hit genes were identified using a FDR of 0.05. Gene scores were normalized by the standard deviation of negative controls genes for each genome-wide sublibrary. Hits were then combined and gene set enrichment analysis (GSEA) was performed for T22 positive and negative bins after filtering for mitochondrial genes using ENRICHR ^90–92^.

### Pairwise analysis secondary screens

After data processing as described above, normalized hits files for re-test screens were compared using custom python scripts to calculate Pearson’s correlation coefficient and generate lists of genes that were unique to each screen in the comparison using a gene score of ±5. We then performed GSEA using ENRICHR on these unique gene sets in order to label categories of genes. Venn diagrams of overlapping hits were generated using a custom python script and the list of overlapping genes processed for GSEA using ENRICHR, after filtering for mitochondrial genes.

## ACKNOWLEDGEMENTS

We would like to thank Dr. Rakez Kayed for tau oligomer specific antibodies. We would like to acknowledge Drs. Carlo Condello, Parker Grosjean, Mor Alkaslasi, Chaitali Anand, Shiaoching Gong, as well as Ian Steele and Molly O’Brien for support and advice. We would like to acknowledge E. Chow (UCSF Center for Advanced Technology) for support with next-generation sequencing; S. Elmes (UCSF Laboratory for Cell Analysis) for support with FACS, Grant P30CA082103. We would like to thank all members of the Kampmann lab for helpful advice and technical support. A.J.S was supported by the Rainwater Charitable Foundation, NIH F32 AG063487, and NIH K99 AG080116-01. M.K. was supported by the Rainwater Charitable Foundation/Tau consortium, the Chan Zuckerberg Initiative Ben Barres Early Career Acceleration Award, the Innovative Genomics Institute and NIH grants R01 AG062359, R01 AG082141, U54 NS100717, U54 NS123746. D.E. and R.A. were supported by NIH U24AG072458 and HHMI. J.E.G was supported by the Tau Consortium. E.C.C was supported by NIH grant F32AG076281. J.G. and V.L.L were supported by NIH grant U54 AI1707922. K.J.L. was supported by NIA U19AG060909.

## AUTHOR CONTRIBUTIONS

A.J.S. and M.K. conceived the project. A.J.S., N.A., designed and performed experiments. A.J.S, J.M., D.L.S, designed and performed mass spectrometry experiments. A.J.S, M.K., G.R., C.P.B, D.G., designed and performed CRISPR-screening experiments. A.J.S., M.K., C.P.B, L.G., designed and performed mouse experiments. A.J.S., M.K., R.B., V.L.L., J.D.G, designed and performed *in vitro* ubiquitination experiments. A.J.S, M.K., E.T., D.R.S, designed and performed negative stain EM experiments. A.J.S, M.K., G.D., E.M., J.J., designed and performed iNeuron fractionation and qPCR experiments. A.J.S., H.P., N.A., G.R., E.C.C., C.P.B., J.M., R.A.,D.E.,R.T.,N.M.K.,J.W.,D.S., L.G., R.E.L., D.R.S, J.E.G, M.K. developed experimental protocols, tools, and reagents or analyzed data. A.J.S. and M.K. wrote the manuscript.

## DECLARATION OF INTERESTS

M. K. is an inventor on US Patent 11,254,933 related to CRISPRi and CRISPRa screening, a co-scientific founder of Montara Therapeutics and serves on the Scientific Advisory Boards of Engine Biosciences, Alector, and Montara Therapeutics, and is an advisor to Modulo Bio and Theseus Therapies. The other authors declare no competing interests.

---

**Figure S1:**
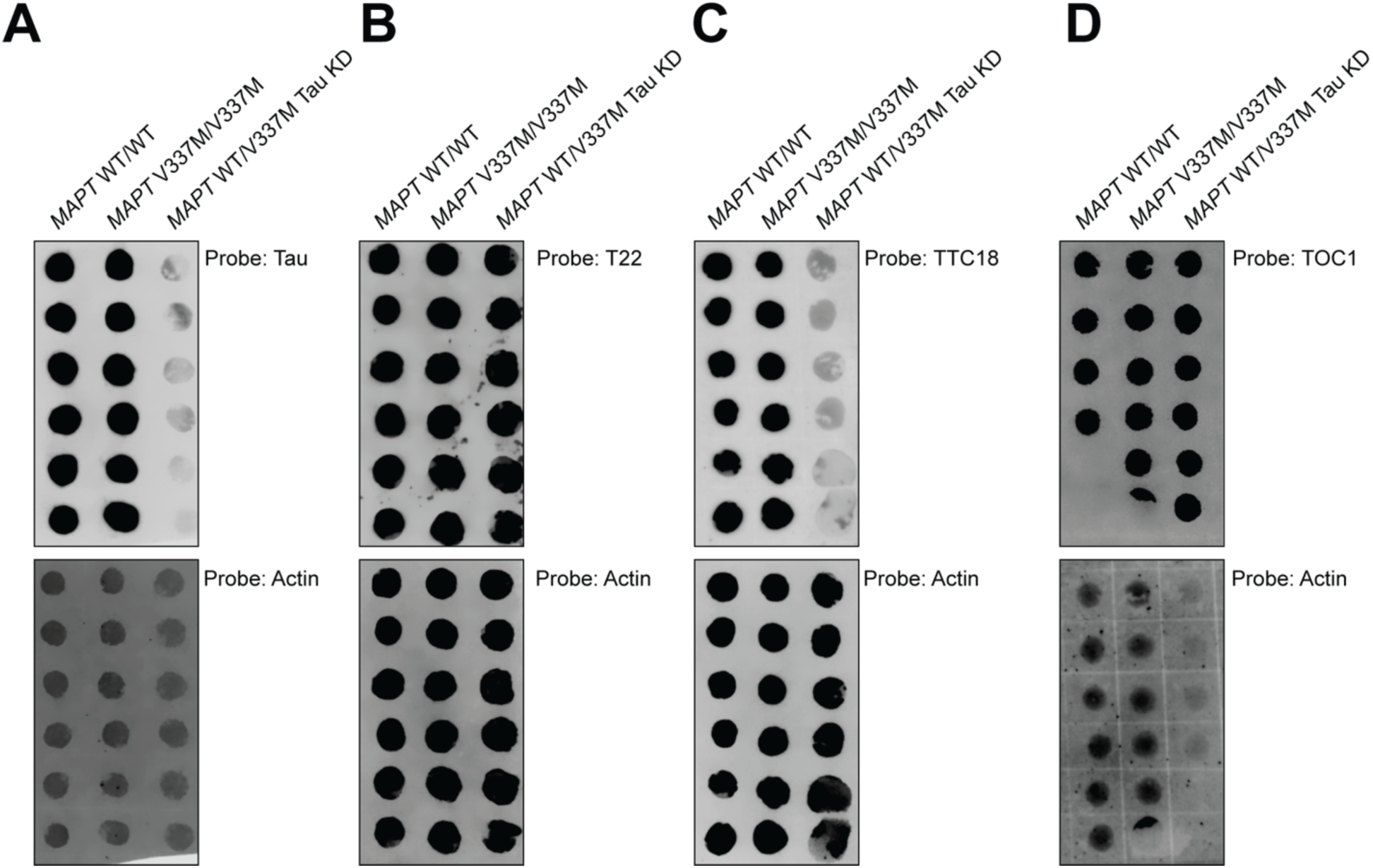
Dot blots used for quantitation in. Figure 1C. Top: Blots probed with either (A) Tau13 (B) T22 (C) TTC18 or (D) TOC1. Below: Blots probed with Actin as a loading control.

**Figure S2:**
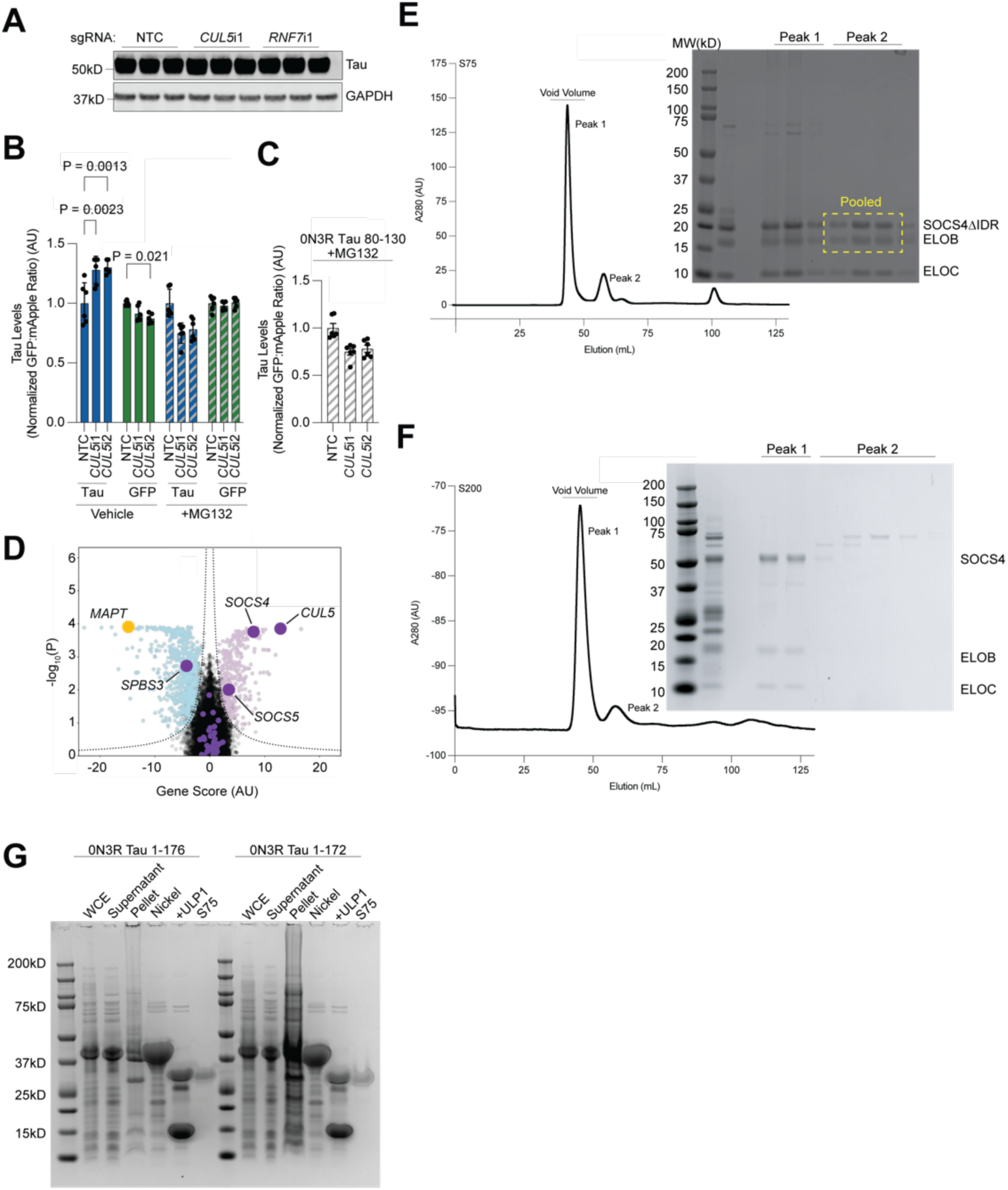
Related to Figure 4 and 7. **(A)** Western blots used in Figure 4B. **(B)** *CUL5* knockdown increases tau levels (blue), but does not increase GFP levels (b*ottom*). Six biological replicates were used per sample. This effect is rescued by treatment with the proteasome inhibitor MG132 (grey patterned bars). **(C)** Treatment of cells expression tau 80-130 with or without *CUL5* knockdown in the presence of the proteasome inhibitor MG132 shows a rescue of *CUL5* KD as compared to vehicle. Six biological replicates were used per sample. For all applicable subpanels, one-way ANOVA was used for statistical analysis. P-values of >0.05 are not shown. Error bars are ±standard deviation. **(D)** Volcano plot of hits from the genome-wide CRISPRi screen for modifiers of tau oligomer levels as in Figure 2, but with all known CRL5 adaptors colored in purple. Only *SOCS4* and, to a lesser degree, *SOCS5* knockdown increase tau oligomer levels. *CUL5* is labeled for reference. **(E)** Chromatogram trace of SOCS4ΔIDR-ELOBC (*left*) and SDS-PAGE of selected peaks (*right*). **(F)** Chromatogram trace of SOCS4 full length (*left*) and SDS-PAGE of selected peaks (*right*). **(G)** Purification of tau 25kD fragments.

**Figure S3:**
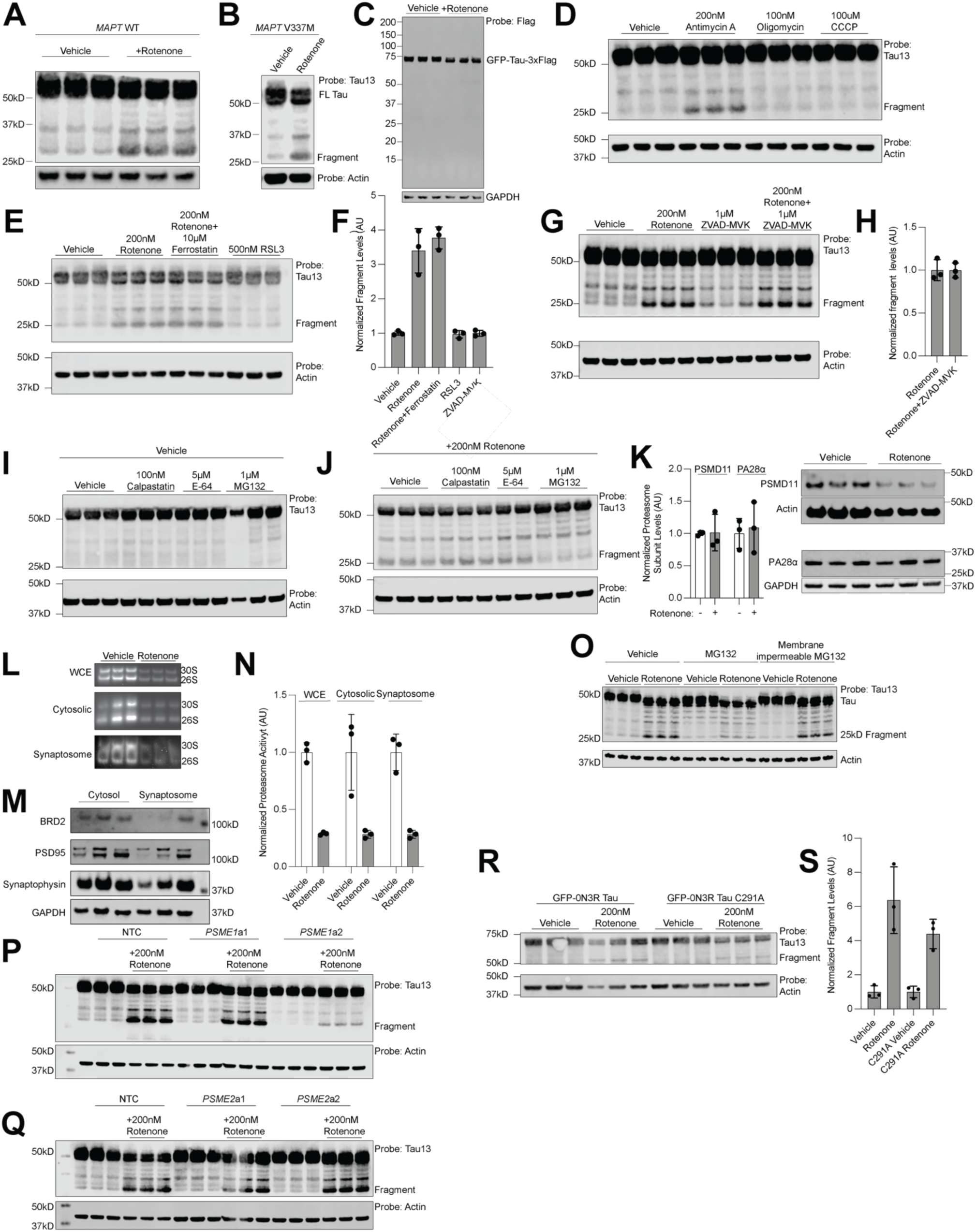
Rotenone treatment leads to 25kD fragment formation. **(A**) Biological triplicate wells of *MAPT* WT neurons treated with vehicle (DMSO) or 200nM rotenone for 24 hours. **(B)** Figure 6C from the main text for comparison with the MAPT V337M tau line. **(C)** Neurons expressing a GFP-0N3R Tau-3xFlag construct treated with vehicle or 200nM rotenone for 24 hours. Probe with an anti-Flag antibody shows no low molecular weight tau fragments. **(D)** Inhibition of ETC complex III, but not complex V or uncoupling of the mitochondrial protein gradient promotes fragment formation. Western blot of vehicle treated, antimycin A treated (complex III inhibitor), oligomycin (complex V inhibitor) or CCCP (protein gradient uncoupler). **(E-H)** Apoptosis and ferroptosis do not control 25kD fragment formation. **(E)** Western blot of rotenone-treated neurons with the ferroptosis inhibitor ferrostatin, or ferroptosis promoting molecule RSL3 reveals no changes to tau 25kD fragment. **(F)** Quantitation of data in (E). **(G)**Western blot of rotenone-treated neurons with the pan-caspase inhibitor ZVAD-MVAK reveals no changes to tau 25kD fragment. **(H)** Quantitation of data in (C). **(I)-(J)** Proteasome inhibition decreases 25kD fragment formation. **(I)** Western blot of neurons with the cathepsin inhibitor E64, calpain inhibitor calpastatin, and proteasome inhibitor MG132. **(J)** Western blot of rotenone treated neurons with the cathepsin inhibitor E64, calpain inhibitor calpastatin, and proteasome inhibitor MG132. **(K)** Levels of PSMD11 and PA28⍺ do not change upon rotenone treatment. Quantitation (*left*) of western blots (*right*). **(L-M)** Synaptic proteasome activity is decreased to the same extent as the entire proteasome pool. **(L)** Proteasome activity assay of proteasomes derived from whole cell lysate or the cytosolic and synaptosome fraction respectively. **(N)** Quantitation of activity from gels in (L)**. (N)** Synaptosome prep reveals depletion of nuclear proteins (BRD2) and enrichment of synaptic proteins. **(O)** Treatment with membrane-impermeable proteasome inhibitor does not affect 25kD fragment formation. **(P-Q)** Gels quantified in Figure 6 **(P)** CRISPRa of PSME1. **(Q)** CRISPRa of PSME2. **(R-S)** Mutation of Tau C291 does not affect fragment formation. **(R)** Western blot of neurons transduced with GFP-tau or GFP-tau C291A treated with rotenone or vehicle. **(S).** Quantitation of data in (R). All samples are the average of three biological replicates, error bars are ±standard deviation.

**Figure S4:**
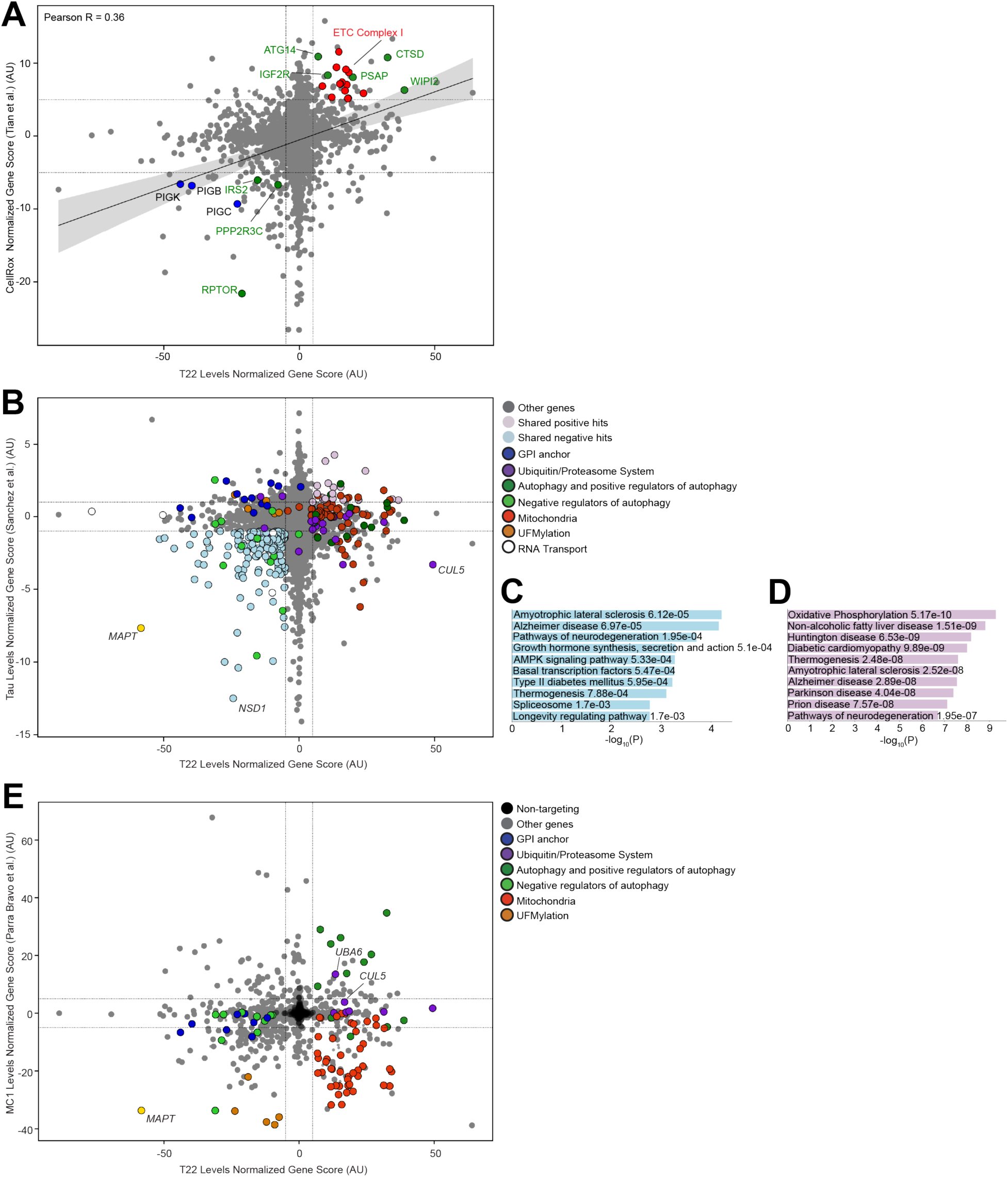
Comparison of screens in this work with others in the literature. **(A)** Shared positive and negative hits genes of Cell Rox (Tian et al.^65^)and T22 screens (this work) were subjected to gene set enrichment analysis. Significant shared KEGG Pathways are labelled above. Red: ETC Complex I, Blue: GPI-anchor biosynthesis, Green: mTOR signaling and autophagy. Other genes involved in oxidative stress are labeled in black. **(B)** Comparison of primary screen with screen for tau levels (Sanchez et al^74^). **(C)** and **(D**) KEGG pathway analysis of shared genes knockdown of which decrease (C) or increase (D) T22 levels (this work) or tau levels (Sanchez et al.). **(E)** Comparison of primary screen with screen for seeding induced tau aggregation (Parra Bravo et al.^76^).

**Figure S5:**
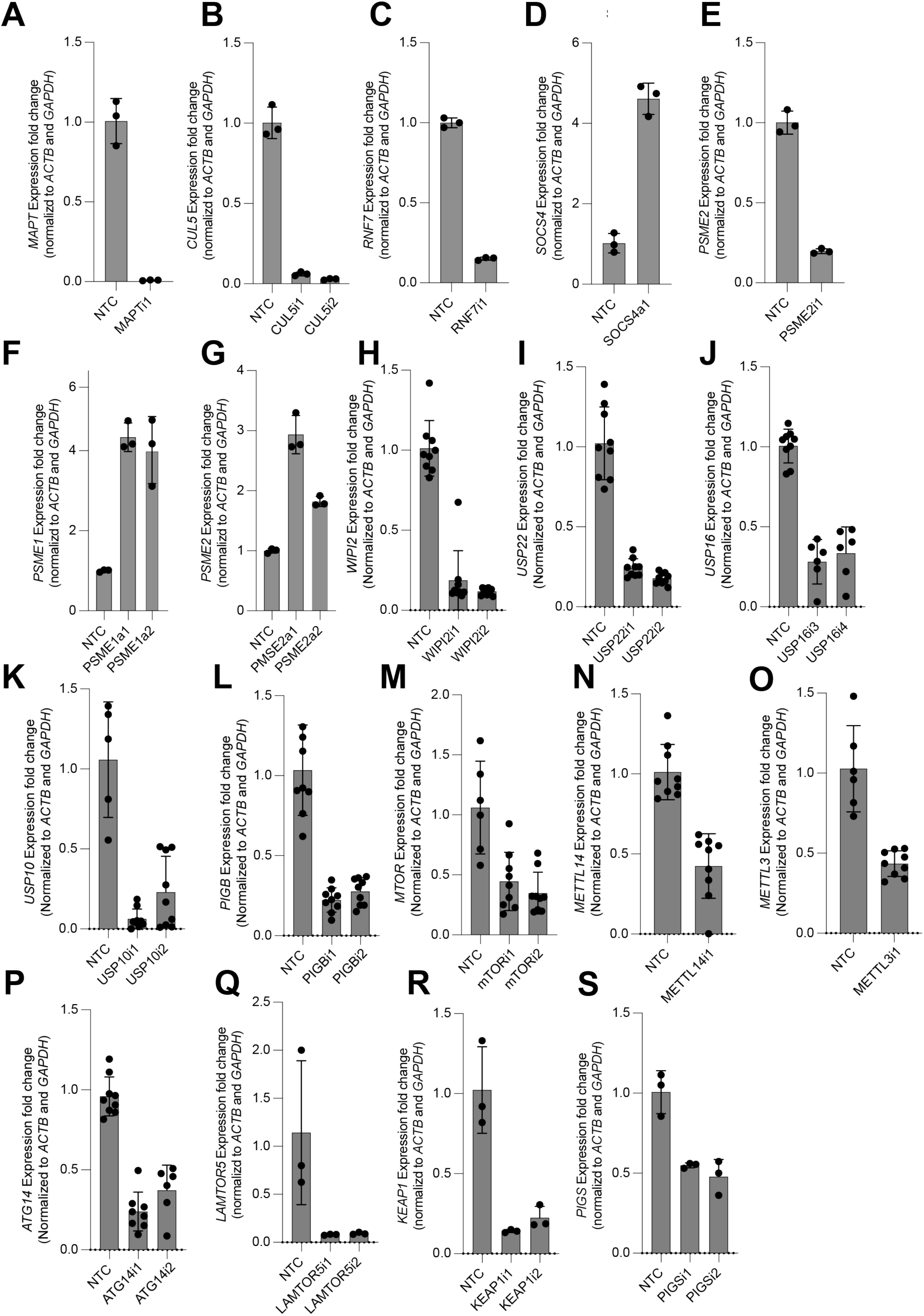
qPCR-based quantification effect of sgRNAs used in this study on expression of the targeted genes. **(A)** sgRNAs targeting *MAPT* in CRISPRi. **(B)** sgRNAs targeting *CUL5* in CRISPRi. **(C)** sgRNAs targeting *RNF7* in CRISPRi. **(D)** sgRNAs targeting *SOCS4* in CRISPRa. **(E)** sgRNAs targeting *PSME2* in CRISPRi. **(F)** sgRNAs targeting *PSME1* in CRISPRa. **(G)** sgRNAs targeting *PSME2* in CRISPRa. **(H)** sgRNAs targeting *WIPI2* in CRISPRi. **(I)** sgRNAs targeting *USP22* in CRISPRi. **(J)** sgRNAs targeting *USP16* in CRISPRi. **(K)** sgRNAs targeting *USP10* in CRISPRi. **(L)** sgRNAs targeting *PIGB* in CRISPRi. **(M)** sgRNAs targeting *MTOR* in CRISPRi. **(N)** sgRNAs targeting *METTL14* in CRISPRi. **(O)** sgRNAs targeting *METTL3* in CRISPRi. **(P)** sgRNAs targeting *ATG14* in CRISPRi. **(Q)** sgRNAs targeting *LAMTOR5* in CRISPRi. **(R)** sgRNAs targeting *KEAP1* in CRISPRi. **(S)** sgRNAs targeting *PIGS* in CRISPRi. (A)-(G), (Q)-(R) Average of three biological replicates, (H)-(P), Average of nine biological replicates. For (K), (M), (O), NTCs are average of six biological replicates. For all samples, error bars are ±standard deviation.

**Figure S6:**
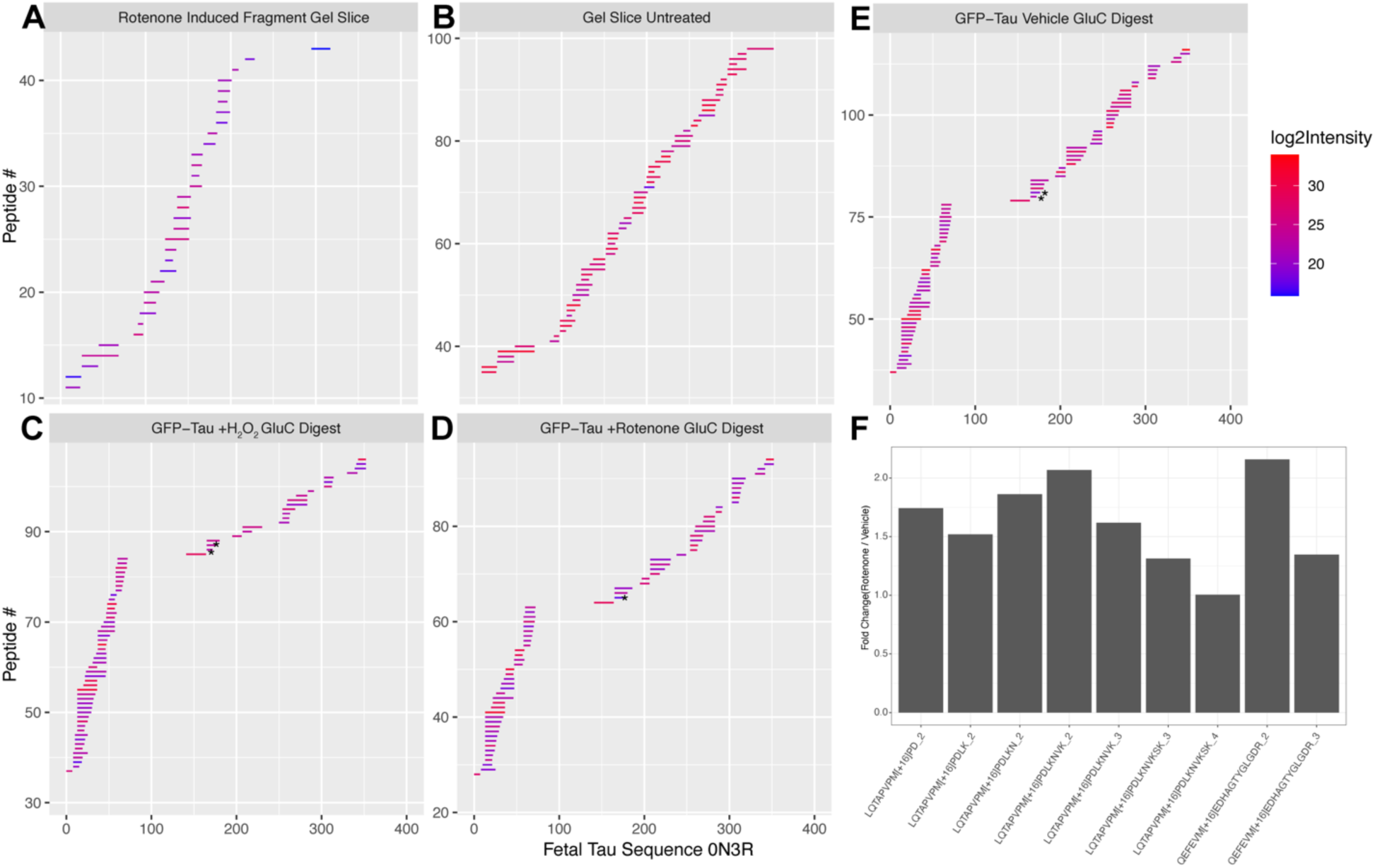
Plots of peptides and their intensities from mass spectrometry experiments. **(A)** Rotenone induced fragment excised from SDS/PAGE gel. **(B)** Full length tau excised from SDS/PAGE gel **(C)** Purified tau peptides from hydrogen peroxide treated neurons digested with GluC. **(D)** Purified tau peptides from 200nM rotenone treated neurons digested with GluC. **(E)** Purified tau peptides from vehicle treated neurons digested with GluC. Stars denote neo-tryptic termini. **(F)** Fold change intensities of oxidized methionine peptides measure by mass spectrometry. Average of four replicates for each peptide.

**Supplemental Table 1 – qPCR primers and sgRNA protospacer sequences used in this study.**

**Supplemental Table 2 – Screen results for primary and secondary screens. Each screen is in a separate tab.**

**Supplemental Table 3 – All hit genes for primary and secondary screens. Each screen is in a separate tab.**

**Supplemental Table 4 – sgRNA protospacers sequences for secondary library**

**Supplemental Table 5 – All identified ubiquitination sites for *in vitro* ubiquitination. Each substrate is in a separate tab.**

**Supplemental Table 6 – All correlation scores calculated from the SEA-AD dataset.**

## Notes

### Summary of Updates

Additional data analysis from human post-mortem brains. Typos fixed.

## REFERENCES

1. Chang, C.W., Shao, E., and Mucke, L. (2021). Tau: Enabler of diverse brain disorders and target of rapidly evolving therapeutic strategies. Science 371. 10.1126/science.abb8255.

2. Wang, Y., and Mandelkow, E. (2016). Tau in physiology and pathology. Nature reviews. Neuroscience 17, 5–21. 10.1038/nrn.2015.1.

3. Tracy, T.E., and Gan, L. (2018). Tau-mediated synaptic and neuronal dysfunction in neurodegenerative disease. Curr Opin Neurobiol 51, 134–138. 10.1016/j.conb.2018.04.027.

4. Hutton, M., Lendon, C.L., Rizzu, P., Baker, M., Froelich, S., Houlden, H., Pickering-Brown, S., Chakraverty, S., Isaacs, A., Grover, A., et al. (1998). Association of missense and 5’-splice-site mutations in tau with the inherited dementia FTDP-17. Nature 393, 702–705. 10.1038/31508.

5. Poorkaj, P., Grossman, M., Steinbart, E., Payami, H., Sadovnick, A., Nochlin, D., Tabira, T., Trojanowski, J.Q., Borson, S., Galasko, D., et al. (2001). Frequency of tau gene mutations in familial and sporadic cases of non-Alzheimer dementia. Arch Neurol 58, 383–387. 10.1001/archneur.58.3.383.

6. van Swieten, J., and Spillantini, M.G. (2007). Hereditary frontotemporal dementia caused by Tau gene mutations. Brain Pathol 17, 63–73. 10.1111/j.1750-3639.2007.00052.x.

7. Goedert, M., and Spillantini, M.G. (2011). Pathogenesis of the tauopathies. Journal of Molecular Neuroscience 45, 425–431. 10.1007/s12031-011-9593-4.

8. Braak, H., and Braak, E. (1991). Neuropathological stageing of Alzheimer-related changes. Acta Neuropathol 82, 239–259. 10.1007/BF00308809.

9. Leng, K., Li, E., Eser, R., Piergies, A., Sit, R., Tan, M., Neff, N., Li, S.H., Rodriguez, R.D., Suemoto, C.K., et al. (2021). Molecular characterization of selectively vulnerable neurons in Alzheimer’s disease. Nat Neurosci 24, 276–287. 10.1038/s41593-020-00764-7.

10. Schö, M., Lockhart, S.N., Schonhaut, D.R., Schwimmer, H.D., Rabinovici, G.D., Correspondence, W.J.J., Schö Ll, M., O ’neil, J.P., Janabi, M., Ossenkoppele, R., et al. (2016). PET Imaging of Tau Deposition in the Aging Human Brain. 971–982. 10.1016/j.neuron.2016.01.028.

11. Seeley, W.W., Crawford, R.K., Zhou, J., Miller, B.L., and Greicius, M.D. (2009). Neurodegenerative diseases target large-scale human brain networks. Neuron 62, 42–52. 10.1016/j.neuron.2009.03.024.

12. Rexach, J.E., Cheng, Y., Chen, L., Polioudakis, D., Lin, L.-C., Mitri, V., Elkins, A., Han, X., Yamakawa, M., Yin, A., et al. (2024). Cross-disorder and disease-specific pathways in dementia revealed by single-cell genomics. Cell 187, 5753–5774.e28. 10.1016/j.cell.2024.08.019.

13. Shi, Y., Zhang, W., Yang, Y., Murzin, A.G., Falcon, B., Kotecha, A., van Beers, M., Tarutani, A., Kametani, F., Garringer, H.J., et al. (2021). Structure-based classification of tauopathies. Nature 598, 359–363. 10.1038/s41586-021-03911-7.

14. Arakhamia, T., Lee, C.E., Carlomagno, Y., Seyfried, N.T., Petrucelli, L., Fitzpatrick, A.W.P., Arakhamia, T., Lee, C.E., Carlomagno, Y., Duong, D.M., et al. (2020). Posttranslational Modifications Mediate the Structural Diversity of Tauopathy Strains. Cell 180, 1–12. 10.1016/j.cell.2020.01.027.

15. Wesseling, H., Mair, W., Kumar, M., Schlaffner, C.N., Tang, S., Beerepoot, P., Fatou, B., Guise, A.J., Cheng, L., Takeda, S., et al. (2020). Tau PTM Profiles Identify Patient Heterogeneity and Stages of Alzheimer’s Disease. Cell 183, 1699–1713.e13. 10.1016/j.cell.2020.10.029.

16. Bellenguez, C., Küçükali, F., Jansen, I.E., Kleineidam, L., Moreno-Grau, S., Amin, N., Naj, A.C., Campos-Martin, R., Grenier-Boley, B., Andrade, V., et al. (2022). New insights into the genetic etiology of Alzheimer’s disease and related dementias. Nat Genet 54, 412–436. 10.1038/s41588-022-01024-z.

17. Wightman, D.P., Jansen, I.E., Savage, J.E., Shadrin, A.A., Bahrami, S., Holland, D., Rongve, A., Børte, S., Winsvold, B.S., Drange, O.K., et al. (2021). A genome-wide association study with 1,126,563 individuals identifies new risk loci for Alzheimer’s disease. Nat Genet 53, 1276–1282. 10.1038/s41588-021-00921-z.

18. Jansen, I.E., Savage, J.E., Watanabe, K., Bryois, J., Williams, D.M., Steinberg, S., Sealock, J., Karlsson, I.K., Hägg, S., Athanasiu, L., et al. (2019). Genome-wide meta-analysis identifies new loci and functional pathways influencing Alzheimer’s disease risk. Nat Genet 51, 404–413. 10.1038/s41588-018-0311-9.

19. Andrews, S.J., Fulton-Howard, B., and Goate, A. (2020). Interpretation of risk loci from genome-wide association studies of Alzheimer’s disease. Lancet Neurol 19, 326–335. 10.1016/S1474-4422(19)30435-1.

20. Mathys, H., Davila-Velderrain, J., Peng, Z., Gao, F., Mohammadi, S., Young, J.Z., Menon, M., He, L., Abdurrob, F., Jiang, X., et al. (2019). Single-cell transcriptomic analysis of Alzheimer’s disease. Nature 570, 332–337. 10.1038/s41586-019-1195-2.

21. Otero-Garcia, M., Mahajani, S.U., Wakhloo, D., Tang, W., Xue, Y.-Q., Morabito, S., Pan, J., Oberhauser, J., Madira, A.E., Shakouri, T., et al. (2022). Molecular signatures underlying neurofibrillary tangle susceptibility in Alzheimer’s disease. Neuron 110, 2929–2948.e8. 10.1016/j.neuron.2022.06.021.

22. Tian, R., Gachechiladze, M.A., Ludwig, C.H., Laurie, M.T., Hong, J.Y., Nathaniel, D., Prabhu, A.V., Fernandopulle, M.S., Patel, R., Abshari, M., et al. (2019). CRISPR Interference-Based Platform for Multimodal Genetic Screens in Human iPSC-Derived Neurons. Neuron 104, 239–255.e12. 10.1016/j.neuron.2019.07.014.

23. Hong, M. (1998). Mutation-Specific Functional Impairments in Distinct Tau Isoforms of Hereditary FTDP-17. Science 282, 1914–1917. 10.1126/science.282.5395.1914.

24. Spina, S., Schonhaut, D.R., Boeve, B.F., Seeley, W.W., Ossenkoppele, R., O’Neil, J.P., Lazaris, A., Rosen, H.J., Boxer, A.L., Perry, D.C., et al. (2017). Frontotemporal dementia with the V337M MAPT mutation: Tau-PET and pathology correlations. Neurology 88, 758– 766. 10.1212/WNL.0000000000003636.

25. Qi, C., Lövestam, S., Murzin, A.G., Peak-Chew, S., Franco, C., Bogdani, M., Latimer, C., Murrell, J.R., Cullinane, P.W., Jaunmuktane, Z., et al. (2024). Tau filaments with the Alzheimer fold in cases with MAPT mutations V337M and R406W. bioRxiv, 2024.04.29.591661. 10.1101/2024.04.29.591661.

26. Snellman, A., Lantero-Rodriguez, J., Emeršič, A., Vrillon, A., Karikari, T.K., Ashton, N.J., Gregorič Kramberger, M., Čučnik, S., Paquet, C., Rot, U., et al. (2022). N-terminal and mid-region tau fragments as fluid biomarkers in neurological diseases. Brain 145, 2834–2848. 10.1093/brain/awab481.

27. Lantero-Rodriguez, J., Salvadó, G., Snellman, A., Montoliu-Gaya, L., Brum, W.S., Benedet, A.L., Mattsson-Carlgren, N., Tideman, P., Janelidze, S., Palmqvist, S., et al. (2024). Plasma N-terminal containing tau fragments (NTA-tau): a biomarker of tau deposition in Alzheimer’s Disease. Mol Neurodegeneration 19, 1–22. 10.1186/s13024-024-00707-x.

28. Lantero-Rodriguez, J., Tissot, C., Snellman, A., Servaes, S., Benedet, A.L., Rahmouni, N., Montoliu-Gaya, L., Therriault, J., Brum, W.S., Stevenson, J., et al. (2023). Plasma and CSF concentrations of N-terminal tau fragments associate with in vivo neurofibrillary tangle burden. Alzheimers Dement 19, 5343–5354. 10.1002/alz.13119.

29. Fá, M., Puzzo, D., Piacentini, R., Staniszewski, A., Zhang, H., Baltrons, M.A., Li Puma, D.D., Chatterjee, I., Li, J., Saeed, F., et al. (2016). Extracellular Tau Oligomers Produce An Immediate Impairment of LTP and Memory. Scientific Reports 6, 1–15. 10.1038/srep19393.

30. Gyparaki, M.T., Arab, A., Sorokina, E.M., Santiago-Ruiz, A.N., Bohrer, C.H., Xiao, J., and Lakadamyali, M. (2021). Tau forms oligomeric complexes on microtubules that are distinct from tau aggregates. Proc Natl Acad Sci U S A 118, e2021461118. 10.1073/pnas.2021461118.

31. Lasagna-Reeves, C.A., Castillo-Carranza, D.L., Sengupta, U., Sarmiento, J., Troncoso, J., Jackson, G.R., and Kayed, R. (2012). Identification of oligomers at early stages of tau aggregation in Alzheimer’s disease. The FASEB Journal 26, 1946–1959. 10.1096/fj.11-199851.

32. Colom-Cadena, M., Davies, C., Sirisi, S., Lee, J.-E., Simzer, E.M., Tzioras, M., Querol-Vilaseca, M., Sánchez-Aced, É., Chang, Y.Y., Holt, K., et al. (2023). Synaptic oligomeric tau in Alzheimer’s disease - A potential culprit in the spread of tau pathology through the brain. Neuron 111, 2170–2183.e6. 10.1016/j.neuron.2023.04.020.

33. Zhang, Z.-Y., Harischandra, D.S., Wang, R., Ghaisas, S., Zhao, J.Y., McMonagle, T.P., Zhu, G., Lacuarta, K.D., Song, J., Trojanowski, J.Q., et al. (2023). TRIM11 protects against tauopathies and is down-regulated in Alzheimer’s disease. Science 381, eadd6696. 10.1126/science.add6696.

34. Abud, E.M., Ramirez, R.N., Martinez, E.S., Healy, L.M., Nguyen, C.H.H., Newman, S.A., Yeromin, A.V., Scarfone, V.M., Marsh, S.E., Fimbres, C., et al. (2017). iPSC-derived human microglia-like cells to study neurological diseases. Neuron 94, 278–293.e9. 10.1016/j.neuron.2017.03.042.

35. Lasagna-Reeves, C.A., Castillo-Carranza, D.L., Sengupta, U., Guerrero-Munoz, M.J., Kiritoshi, T., Neugebauer, V., Jackson, G.R., and Kayed, R. (2012). Alzheimer brain-derived tau oligomers propagate pathology from endogenous tau. Scientific Reports 2. 10.1038/srep00700.

36. Horlbeck, M.A., Gilbert, L.A., Villalta, J.E., Adamson, B., Pak, R.A., Chen, Y., Fields, A.P., Park, C.Y., Corn, J.E., Kampmann, M., et al. (2016). Compact and highly active next-generation libraries for CRISPR-mediated gene repression and activation. eLife 5, 1–20. 10.7554/eLife.19760.

37. Caballero, B., Bourdenx, M., Luengo, E., Diaz, A., Sohn, P.D., Chen, X., Wang, C., Juste, Y.R., Wegmann, S., Patel, B., et al. (2021). Acetylated tau inhibits chaperone-mediated autophagy and promotes tau pathology propagation in mice. Nat Commun 12, 2238. 10.1038/s41467-021-22501-9.

38. Djajadikerta, A., Keshri, S., Pavel, M., Prestil, R., Ryan, L., and Rubinsztein, D.C. (2020). Autophagy Induction as a Therapeutic Strategy for Neurodegenerative Diseases. J Mol Biol 432, 2799–2821. 10.1016/j.jmb.2019.12.035.

39. Menzies, F.M., Fleming, A., Caricasole, A., Bento, C.F., Andrews, S.P., Ashkenazi, A., Füllgrabe, J., Jackson, A., Jimenez Sanchez, M., Karabiyik, C., et al. (2017). Autophagy and Neurodegeneration: Pathogenic Mechanisms and Therapeutic Opportunities. Neuron 93, 1015–1034. 10.1016/j.neuron.2017.01.022.

40. Silva, M.C., Nandi, G.A., Tentarelli, S., Gurrell, I.K., Jamier, T., Lucente, D., Dickerson, B.C., Brown, D.G., Brandon, N.J., and Haggarty, S.J. (2020). Prolonged tau clearance and stress vulnerability rescue by pharmacological activation of autophagy in tauopathy neurons. Nat Commun 11, 3258. 10.1038/s41467-020-16984-1.

41. Jiang, L., Lin, W., Zhang, C., Ash, P.E.A., Verma, M., Kwan, J., van Vliet, E., Yang, Z., Cruz, A.L., Boudeau, S., et al. (2021). Interaction of tau with HNRNPA2B1 and N6- methyladenosine RNA mediates the progression of tauopathy. Molecular Cell 81, 4209–4227.e12. 10.1016/j.molcel.2021.07.038.

42. Kanaan, N.M., Cox, K., Alvarez, V.E., Stein, T.D., Poncil, S., and McKee, A.C. (2016). Characterization of Early Pathological Tau Conformations and Phosphorylation in Chronic Traumatic Encephalopathy. J Neuropathol Exp Neurol 75, 19–34. 10.1093/jnen/nlv001.

43. Abskharon, R., Seidler, P.M., Sawaya, M.R., Cascio, D., Yang, T.P., Philipp, S., Williams, C.K., Newell, K.L., Ghetti, B., DeTure, M.A., et al. (2020). Crystal structure of a conformational antibody that binds tau oligomers and inhibits pathological seeding by extracts from donors with Alzheimer’s disease. The Journal of biological chemistry, 1–28. 10.1074/jbc.RA120.013638.

44. Bano, I., Soomro, A.S., Abbas, S.Q., Ahmadi, A., Hassan, S.S.U., Behl, T., and Bungau, S. (2022). A Comprehensive Review of Biological Roles and Interactions of Cullin-5 Protein. ACS Omega 7, 5615–5624. 10.1021/acsomega.1c06890.

45. Gao, F., Fan, Y., Zhou, B., Guo, W., Jiang, X., Shi, J., and Ren, C. (2020). The functions and properties of cullin-5, a potential therapeutic target for cancers. Am J Transl Res 12, 618– 632.

46. Yu, X., Yu, Y., Liu, B., Luo, K., Kong, W., Mao, P., and Yu, X.-F. (2003). Induction of APOBEC3G ubiquitination and degradation by an HIV-1 Vif-Cul5-SCF complex. Science 302, 1056–1060. 10.1126/science.1089591.

47. Xu, P., Liu, Y., Liu, C., Guey, B., Li, L., Melenec, P., Ricci, J., and Ablasser, A. (2024). The CRL5–SPSB3 ubiquitin ligase targets nuclear cGAS for degradation. Nature, 1–7. 10.1038/s41586-024-07112-w.

48. Scott, D.C., Rhee, D.Y., Duda, D.M., Kelsall, I.R., Olszewski, J.L., Paulo, J.A., de Jong, A., Ovaa, H., Alpi, A.F., Harper, J.W., et al. (2016). Two Distinct Types of E3 Ligases Work in Unison to Regulate Substrate Ubiquitylation. Cell 166, 1198–1214.e24. 10.1016/j.cell.2016.07.027.

49. Horn-Ghetko, D., Krist, D.T., Prabu, J.R., Baek, K., Mulder, M.P.C., Klügel, M., Scott, D.C., Ovaa, H., Kleiger, G., and Schulman, B.A. (2021). Ubiquitin ligation to F-box protein targets by SCF-RBR E3-E3 super-assembly. Nature 590, 671–676. 10.1038/s41586-021-03197-9.

50. Kostrhon, S., Prabu, J.R., Baek, K., Horn-Ghetko, D., von Gronau, S., Klügel, M., Basquin, J., Alpi, A.F., and Schulman, B.A. (2021). CUL5-ARIH2 E3-E3 ubiquitin ligase structure reveals cullin-specific NEDD8 activation. Nat Chem Biol 17, 1075–1083. 10.1038/s41589-021-00858-8.

51. Duda, D.M., Borg, L.A., Scott, D.C., Hunt, H.W., Hammel, M., and Schulman, B.A. (2008). Structural insights into NEDD8 activation of cullin-RING ligases: conformational control of conjugation. Cell 134, 995–1006. 10.1016/j.cell.2008.07.022.

52. Baek, K., Scott, D.C., and Schulman, B.A. (2021). NEDD8 and ubiquitin ligation by cullin-RING E3 ligases. Curr Opin Struct Biol 67, 101–109. 10.1016/j.sbi.2020.10.007.

53. Arora, A., Goering, R., Lo, H.-Y.G., and Taliaferro, M.J. (2021). Mechanical Fractionation of Cultured Neuronal Cells into Cell Body and Neurite Fractions. Bio Protoc 11, e4048. 10.21769/BioProtoc.4048.

54. Nadel, C.M., Pokhrel, S., Wucherer, K., Oehler, A., Thwin, A.C., Basu, K., Callahan, M.D., Southworth, D.R., Mordes, D.A., Craik, C.S., et al. (2024). Phosphorylation of tau at a single residue inhibits binding to the E3 ubiquitin ligase, CHIP. Nat Commun 15, 7972. 10.1038/s41467-024-52075-1.

55. Petrucelli, L., Dickson, D., Kehoe, K., Taylor, J., Snyder, H., Grover, A., De Lucia, M., McGowan, E., Lewis, J., Prihar, G., et al. (2004). CHIP and Hsp70 regulate tau ubiquitination, degradation and aggregation. Hum Mol Genet 13, 703–714. 10.1093/hmg/ddh083.

56. Parolini, F., Ataie Kachoie, E., Leo, G., Civiero, L., Bubacco, L., Arrigoni, G., Munari, F., Assfalg, M., D’Onofrio, M., and Capaldi, S. (2023). Site-Specific Ubiquitination of Tau Amyloids Promoted by the E3 Ligase CHIP. Angew Chem Int Ed Engl 62, e202310230. 10.1002/anie.202310230.

57. Bullock, A.N., Rodriguez, M.C., Debreczeni, J.E., Songyang, Z., and Knapp, S. (2007). Structure of the SOCS4-ElonginB/C complex reveals a distinct SOCS box interface and the molecular basis for SOCS-dependent EGFR degradation. Structure 15, 1493–1504. 10.1016/j.str.2007.09.016.

58. Gabitto, M.I., Travaglini, K.J., Rachleff, V.M., Kaplan, E.S., Long, B., Ariza, J., Ding, Y., Mahoney, J.T., Dee, N., Goldy, J., et al. (2023). Integrated multimodal cell atlas of Alzheimer’s disease. Res Sq, rs.3.rs-2921860. 10.21203/rs.3.rs-2921860/v1.

59. Batra, S., Vaquer-Alicea, J., Valdez, C., Taylor, S.P., Manon, V.A., Vega, A.R., Kashmer, O.M., Kolay, S., Lemoff, A., Cairns, N.J., et al. (2025). VCP regulates early tau seed amplification via specific cofactors. Mol Neurodegener 20, 2. 10.1186/s13024-024-00783-z.

60. Saha, I., Yuste-Checa, P., Da Silva Padilha, M., Guo, Q., Körner, R., Holthusen, H., Trinkaus, V.A., Dudanova, I., Fernández-Busnadiego, R., Baumeister, W., et al. (2023). The AAA+ chaperone VCP disaggregates Tau fibrils and generates aggregate seeds in a cellular system. Nat Commun 14, 560. 10.1038/s41467-023-36058-2.

61. Petrucelli, L., Dickson, D., Kehoe, K., Taylor, J., Snyder, H., Grover, A., De Lucia, M., McGowan, E., Lewis, J., Prihar, G., et al. (2004). CHIP and Hsp70 regulate tau ubiquitination, degradation and aggregation. Hum Mol Genet 13, 703–714. 10.1093/hmg/ddh083.

62. Sahara, N., Murayama, M., Mizoroki, T., Urushitani, M., Imai, Y., Takahashi, R., Murata, S., Tanaka, K., and Takashima, A. (2005). In vivo evidence of CHIP up-regulation attenuating tau aggregation. J Neurochem 94, 1254–1263. 10.1111/j.1471-4159.2005.03272.x.

63. Combs, B., Hamel, C., and Kanaan, N.M. (2016). Pathological conformations involving the amino terminus of tau occur early in Alzheimer’s disease and are differentially detected by monoclonal antibodies. Neurobiol Dis 94, 18–31. 10.1016/j.nbd.2016.05.016.

64. Chhatwal, J.P., Schultz, A.P., Dang, Y., Ostaszewski, B., Liu, L., Yang, H.-S., Johnson, K.A., Sperling, R.A., and Selkoe, D.J. (2020). Plasma N-terminal tau fragment levels predict future cognitive decline and neurodegeneration in healthy elderly individuals. Nat Commun 11, 6024. 10.1038/s41467-020-19543-w.

65. Tian, R., Abarientos, A., Hong, J., Hashemi, S.H., Yan, R., Dräger, N., Leng, K., Nalls, M.A., Singleton, A.B., Xu, K., et al. (2021). Genome-wide CRISPRi/a screens in human neurons link lysosomal failure to ferroptosis. Nature Neuroscience 24, 1020–1034. 10.1038/s41593-021-00862-0.

66. David, D.C., Layfield, R., Serpell, L., Narain, Y., Goedert, M., and Spillantini, M.G. (2002). Proteasomal degradation of tau protein. J Neurochem 83, 176–185. 10.1046/j.1471-4159.2002.01137.x.

67. Quinn, J.P., Corbett, N.J., Kellett, K.A.B., and Hooper, N.M. (2018). Tau Proteolysis in the Pathogenesis of Tauopathies: Neurotoxic Fragments and Novel Biomarkers. Journal of Alzheimer’s disease : JAD 63, 13–33. 10.3233/JAD-170959.

68. Sampognaro, P.J., Arya, S., Knudsen, G.M., Gunderson, E.L., Sandoval-Perez, A., Hodul, M., Bowles, K., Craik, C.S., Jacobson, M.P., and Kao, A.W. (2023). Mutations in α-synuclein, TDP-43 and tau prolong protein half-life through diminished degradation by lysosomal proteases. Mol Neurodegener *18*, 29. 10.1186/s13024-023-00621-8.

69. Ukmar-Godec, T., Fang, P., Ibáñez de Opakua, A., Henneberg, F., Godec, A., Pan, K.T., Cima-Omori, M.S., Chari, A., Mandelkow, E., Urlaub, H., et al. (2020). Proteasomal degradation of the intrinsically disordered protein tau at single-residue resolution. Science Advances 6, 1–13. 10.1126/sciadv.aba3916.

70. Kors, S., Geijtenbeek, K., Reits, E., and Schipper-Krom, S. (2019). Regulation of Proteasome Activity by (Post-)transcriptional Mechanisms. Front Mol Biosci 6, 48. 10.3389/fmolb.2019.00048.

71. Rechsteiner, M., and Hill, C.P. (2005). Mobilizing the proteolytic machine: cell biological roles of proteasome activators and inhibitors. Trends Cell Biol 15, 27–33. 10.1016/j.tcb.2004.11.003.

72. Duan, L., Hu, M., Tamm, J.A., Grinberg, Y.Y., Shen, F., Chai, Y., Xi, H., Gibilisco, L., Ravikumar, B., Gautam, V., et al. (2021). Arrayed CRISPR reveals genetic regulators of tau aggregation, autophagy and mitochondria in Alzheimer’s disease model. Sci Rep 11, 2879. 10.1038/s41598-021-82658-7.

73. Kim, J., de Haro, M., Al-Ramahi, I., Garaicoechea, L.L., Jeong, H.-H., Sonn, J.Y., Tadros, B., Liu, Z., Botas, J., and Zoghbi, H.Y. (2023). Evolutionarily conserved regulators of tau identify targets for new therapies. Neuron 111, 824–838.e7. 10.1016/j.neuron.2022.12.012.

74. Sanchez, C.G., Acker, C.M., Gray, A., Varadarajan, M., Song, C., Cochran, N.R., Paula, S., Lindeman, A., An, S., McAllister, G., et al. (2021). Genome-wide CRISPR screen identifies protein pathways modulating tau protein levels in neurons. Communications Biology 4, 1–14. 10.1038/s42003-021-02272-1.

75. Yan, Y., Wang, X., Chaput, D., Shin, M.-K., Koh, Y., Gan, L., Pieper, A.A., Woo, J.-A.A., and Kang, D.E. (2022). X-linked ubiquitin-specific peptidase 11 increases tauopathy vulnerability in women. Cell 185, 3913–3930.e19. 10.1016/j.cell.2022.09.002.

76. Parra Bravo, C., Giani, A.M., Madero-Perez, J., Zhao, Z., Wan, Y., Samelson, A.J., Wong, M.Y., Evangelisti, A., Cordes, E., Fan, L., et al. (2024). Human iPSC 4R tauopathy model uncovers modifiers of tau propagation. Cell, S0092–8674(24)00306-4. 10.1016/j.cell.2024.03.015.

77. Cicognola, C., Satir, T.M., Brinkmalm, G., Matečko-Burmann, I., Agholme, L., Bergström, P., Becker, B., Zetterberg, H., Blennow, K., and Höglund, K. (2020). Tauopathy-Associated Tau Fragment Ending at Amino Acid 224 Is Generated by Calpain-2 Cleavage. Journal of Alzheimer’s Disease 74, 1143–1156. 10.3233/JAD-191130.

78. Ramani, B., Rose, I.V.L., Teyssier, N., Pan, A., Danner-Bocks, S., Sanghal, T., Yadanar, L., Tian, R., Ma, K., Palop, J.J., et al. (2025). CRISPR screening by AAV episome-sequencing (CrAAVe-seq): a scalable cell-type-specific in vivo platform uncovers neuronal essential genes. Nat Neurosci. 10.1038/s41593-025-02043-9.

79. Sohn, P.D., Huang, C.T., Yan, R., Xu, K., Kosik, K.S., Gan, L., Sohn, P.D., Huang, C.T., Yan, R., Fan, L., et al. (2019). Pathogenic Tau Impairs Axon Initial Segment Plasticity and Excitability Homeostasis Article Pathogenic Tau Impairs Axon Initial Segment Plasticity and Excitability Homeostasis. 1–13. 10.1016/j.neuron.2019.08.008.

80. Deshar, R., Moon, S., Yoo, W., Cho, E.-B., Yoon, S.K., and Yoon, J.-B. (2016). RNF167 targets Arl8B for degradation to regulate lysosome positioning and endocytic trafficking. FEBS J 283, 4583–4599. 10.1111/febs.13947.

81. Wang, C., Ward, M.E., Chen, R., Liu, K., Tracy, T.E., Chen, X., Xie, M., Sohn, P.D., Ludwig, C., Meyer-Franke, A., et al. (2017). Scalable Production of iPSC-Derived Human Neurons to Identify Tau-Lowering Compounds by High-Content Screening. Stem Cell Reports 9, 1221–1233. 10.1016/j.stemcr.2017.08.019.

82. Gilbert, L.A., Horlbeck, M.A., Adamson, B., Villalta, J.E., Chen, Y., Whitehead, E.H., Guimaraes, C., Panning, B., Ploegh, H.L., Bassik, M.C., et al. (2014). Genome-Scale CRISPR-Mediated Control of Gene Repression and Activation. Cell 159, 647–661. 10.1016/j.cell.2014.09.029.

83. Hüttenhain, R., Xu, J., Burton, L.A., Gordon, D.E., Hultquist, J.F., Johnson, J.R., Satkamp, L., Hiatt, J., Rhee, D.Y., Baek, K., et al. (2019). ARIH2 Is a Vif-Dependent Regulator of CUL5-Mediated APOBEC3G Degradation in HIV Infection. Cell Host Microbe 26, 86–99.e7. 10.1016/j.chom.2019.05.008.

84. Lövestam, S., Koh, F.A., van Knippenberg, B., Kotecha, A., Murzin, A.G., Goedert, M., and Scheres, S.H. (2022). Assembly of recombinant tau into filaments identical to those of Alzheimer’s disease and chronic traumatic encephalopathy. eLife 11, e76494. 10.7554/eLife.76494.

85. Montgomery, K.M., Carroll, E.C., Thwin, A.C., Quddus, A.Y., Hodges, P., Southworth, D.R., and Gestwicki, J.E. (2023). Chemical Features of Polyanions Modulate Tau Aggregation and Conformational States. J Am Chem Soc 145, 3926–3936. 10.1021/jacs.2c08004.

86. He, L., Davila-Velderrain, J., Sumida, T.S., Hafler, D.A., Kellis, M., and Kulminski, A.M. (2021). NEBULA is a fast negative binomial mixed model for differential or co-expression analysis of large-scale multi-subject single-cell data. Commun Biol 4, 1–17. 10.1038/s42003-021-02146-6.

87. Büttner, M., Ostner, J., Müller, C.L., Theis, F.J., and Schubert, B. (2021). scCODA is a Bayesian model for compositional single-cell data analysis. Nat Commun 12, 6876. 10.1038/s41467-021-27150-6.

88. Street, K., Risso, D., Fletcher, R.B., Das, D., Ngai, J., Yosef, N., Purdom, E., and Dudoit, S. (2018). Slingshot: cell lineage and pseudotime inference for single-cell transcriptomics. BMC Genomics 19, 1–16. 10.1186/s12864-018-4772-0.

89. Negative Staining and Image Classification - Powerful Tools in Modern Electron Microscopy - PubMed https://pubmed.ncbi.nlm.nih.gov/15103397/.

90. Chen, E.Y., Tan, C.M., Kou, Y., Duan, Q., Wang, Z., Meirelles, G., Clark, N.R., and Ma’ayan, A. (2013). Enrichr: interactive and collaborative HTML5 gene list enrichment analysis tool. BMC Bioinformatics 14, 128. 10.1186/1471-2105-14-128.

91. Kuleshov, M.V., Jones, M.R., Rouillard, A.D., Fernandez, N.F., Duan, Q., Wang, Z., Koplev, S., Jenkins, S.L., Jagodnik, K.M., Lachmann, A., et al. (2016). Enrichr: a comprehensive gene set enrichment analysis web server 2016 update. Nucleic Acids Res 44, W90–97. 10.1093/nar/gkw377.

92. Xie, Z., Bailey, A., Kuleshov, M.V., Clarke, D.J.B., Evangelista, J.E., Jenkins, S.L., Lachmann, A., Wojciechowicz, M.L., Kropiwnicki, E., Jagodnik, K.M., et al. (2021). Gene Set Knowledge Discovery with Enrichr. Curr Protoc 1, e90. 10.1002/cpz1.90.

